# Tailored synthetic sRNAs dynamically tune multilayer genetic circuits

**DOI:** 10.1101/2022.11.09.515794

**Authors:** Ana K. Velazquez Sanchez, Bjarne Klopprogge, Karl-Heinz Zimmermann, Zoya Ignatova

## Abstract

Predictable and controllable tuning of genetic circuits to regulate gene expression, including modulation of existing circuits or constructs without the need for redesign or rebuilding, is a persistent challenge in synthetic biology. Here, we propose a rational design of de novo small RNAs (sRNAs) to dynamically modulate gene expression within a broad range, from high, medium to low repression, and implemented them in *Escherichia coli*. We designed multiple multilayer genetic circuits, in which the variable effector element is a transcription factor (TF) controlling downstream the production of a reporter protein. Our approach harnesses the intrinsic RNA-interference pathway in *E. coli* to exert dynamic and modular control of the multilayer genetic circuits. The sRNAs were designed to target TFs instead of the reporter gene, which allowed for wide range of expression modulation of the reporter protein, including the most difficult to achieve dynamic switch to an OFF state. Out work provides a frame for achieving independent modulation of gene expression, by only including an independent control circuit expressing synthetic sRNAs, without altering the structure of existing genetic circuits.

## Introduction

Synthetic biology (SynBio) aims at introducing de novo functionalities into living organisms by rational design and interconnection of biological parts (e.g. promoters, transcription factors, regulatory proteins, and RNA) from different sources (e.g. bacteria, bacteriophages, yeast, mammalian systems) (Pasotti *et al*, 2013; Cameron *et al*, 2014; Brophy & Voigt, 2014; Lammens *et al*, 2020). Usually, those synthetic constructs are powered primarily by transcription networks, where regulatory or reporter proteins are under the control of promoters, which respond to one or more transcription factors (TFs). However, frequently those systems present variations in rates of transcription and translation in vivo, depending on multiple intrinsic and extrinsic factors (Rudge *et al*, 2016), including growth rate, availability of cellular resources, and experimental environment (Hooshangi *et al*, 2005; Ceroni *et al*, 2018; Kelly *et al*, 2018). Fine-tuning genetic regulatory components for achieving desired functions or outputs is still a persistent challenge in the field (Bartoli *et al*, 2020; Quarton *et al*, 2020; Ceroni *et al*, 2018). Thus, the development of versatile and programmable genetic expression regulators is desirable. Thereby, ligand-inducible (i.e. TF-controllable) genetic constructs must exhibit tight and well-defined ON and OFF functions and a broad dynamic range (Chen *et al*, 2018). The switch of an ON to an OFF state, is referred to as NOT function or gate, and it is often achieved by negatively inducible promoters that repress transcription in presence of a specific ligand, eventually ceasing the production of the final protein output. An example for this is P_tet_ promoter which is constitutively ON, but following TetR binding it is switched off and transcription is shut down. Traditional gene knockout strategies with CRISPR-CS9, or suppressor mutations, enable a selective termination of production by suppressing or turning OFF genes; however, this effect is irreversible, because of the introduced genomic changes.

Harnessing the intrinsic cellular RNA interference (RNAi) machinery (Zamore *et al*, 2000), a targeted knockdown of genes can be achieved (Wiedenheft *et al*, 2012; Dietzl *et al*, 2007; Sui *et al*, 2002). Across all organisms, small noncoding RNAs (sRNAs) are present as trans-acting regulators of translation and stability of messenger RNAs (mRNAs) (Gottesman, 2004). Engineering synthetic molecules that mimic naturally occurring sRNAs to perform various regulatory tasks have expanded the SynBio toolkit (Liang *et al*, 2011; Yoo *et al*, 2013; Doudna & Charpentier, 2014; Mandal *et al*, 2004; Taxman *et al*, 2010). Bacterial sRNAs regulate gene expression both positively and negatively, by modulating the rates of translation initiation and degradation of their target mRNAs (Bandyra *et al*, 2012). In *Escherichia coli*, a large group of negative regulator sRNAs contains a consensus structured scaffold to recruit the RNA chaperone Hfq (Urban & Vogel, 2007), which promotes annealing between sRNAs and target mRNAs and guides RNase E to degrade mRNA (Morita *et al*, 2005). sRNAs negatively regulate translation of their targets by covering either the Shine Dalgarno (SD) sequence or the start codon (AUG) (Papenfort *et al*, 2010), thus, disfavoring 30S ribosomal subunit biding and translation initiation (Bouvier *et al*, 2008).

Here, we propose a rational design of synthetic sRNAs with different strength of repression (low, medium and high) to dynamically modulate gene expression. We explored the utilization of tailored synthetic sRNAs to dynamically control the expression of TFs, which modulate the output of a downstream fluorescent reporter of multilayered genetic circuits. We tested the modulatory effect of the synthetic sRNAs using three different TFs, and show that different dynamic range of repression can be achieved. Our results suggest that gene expression can be dynamically modulated by incorporating synthetic sRNAs to existing genetic constructs, without modifications of the genetic parts (e.g. exchanging promoters) across different conditions (i.e. time of induction, variation of inducer concentration).

## Results

### Design of synthetic sRNAs to achieve different levels of repression

We customized the sRNAs with an antisense sequence for the target mRNA to repress, immediately followed by the Hfq-recruiting motif which is taken from the naturally occurring sRNA MicC (Na *et al*, 2013; Yoo *et al*, 2013) (Figure 1A). We chose different length of the antisense sequence dependent on the desired repression efficiency. The binding energy between the antisense sRNA module (only target-binding sequence) and its cognate mRNA correlates with the repression activity (Na *et al*, 2013), enabling the prediction of the sRNA efficiency. The degree of suppression and its correlation with the binding energy has been empirically determined (Yoo *et al*, 2013), so that sRNAs with a binding energy of -40 to -30 kcal/mol exhibit high translation repression (up to 90%), whereas sRNAs with binding energy of -15 kcal/mol poorly repress translation (∼10%). Based on this, we put thresholds to obtain three different suppression levels: high with binding energy of -40 to -30 kcal/mol, medium with -30 to -20 kcal/mol and low with -20 to - 15 kcal/mol (Figure 1A). To tune the binding energy within each range, we deleted, added, or randomly exchanged nucleotides within the antisense module (Figure S1).

**Figure 1.**
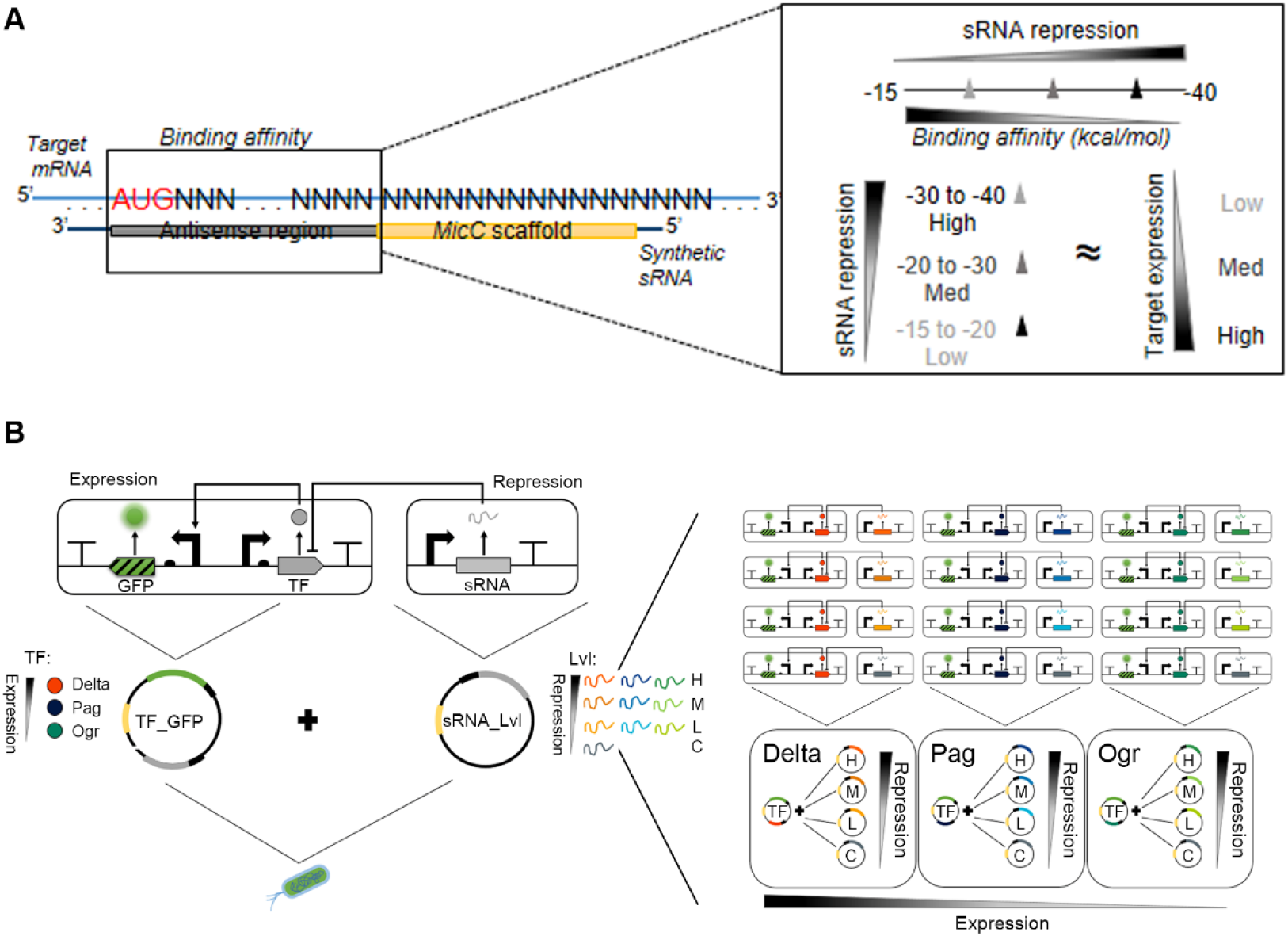
Overview of the engineered modular synthetic sRNAs. **(A)** Schematic of the rational design of synthetic sRNAs. **(B)** Conceptual design of the sRNA tunable multi-layered circuits *in vivo*. Transcription regulation strength: red, TF with high expression (Delta); blue, TF with medium expression (Pag); green, TF with low expression (Ogr). Regulation strength: H, high repression; M, medium repression; L, low repression. C, scrambled sRNA serving as a negative control.

### Engineering multilayered synthetic circuits with different transcriptional activity and protein production yield

Instead of directly repressing the reporter gene in the circuit, we explored the dynamic modulation with synthetic sRNAs by targeting TFs; TFs trigger cascade reactions in circuits. We constructed different multilayer circuits (Figure 1B) and each circuit consists of a fluorescent protein reporter (GFP) and a TF gene (i.e. *δ* TF (Delta) from P4-related retrophage *ϕ*R73 (Slettan *et al*, 1992), Pag TF from PSP3 bacteriophage (Julien & Calendar, 1996), Ogr TF from P2 phage (Slettan *et al*, 1992)). All selected TFs activate the P_F_ promoter, however they differ in their transcription activity and consequently led to different expression yields of the GFP reporter (i.e. strong, moderate, weak) (Figure 2 and S2A,B). The first layer of the multilayer circuits corresponds to the production of a TF under control of the P_BAD_ promoter induced by arabinose; TF expression switches on a second layer of expression of GFP under the P_F_ promoter from the phage P2 and the GFP fluorescence is the output signal of the circuit. The third layer corresponds to post-transcriptional TF regulation or the second layer effector, constituting a synthetic sRNAs targeting the TF mRNA.

**Figure 2.**
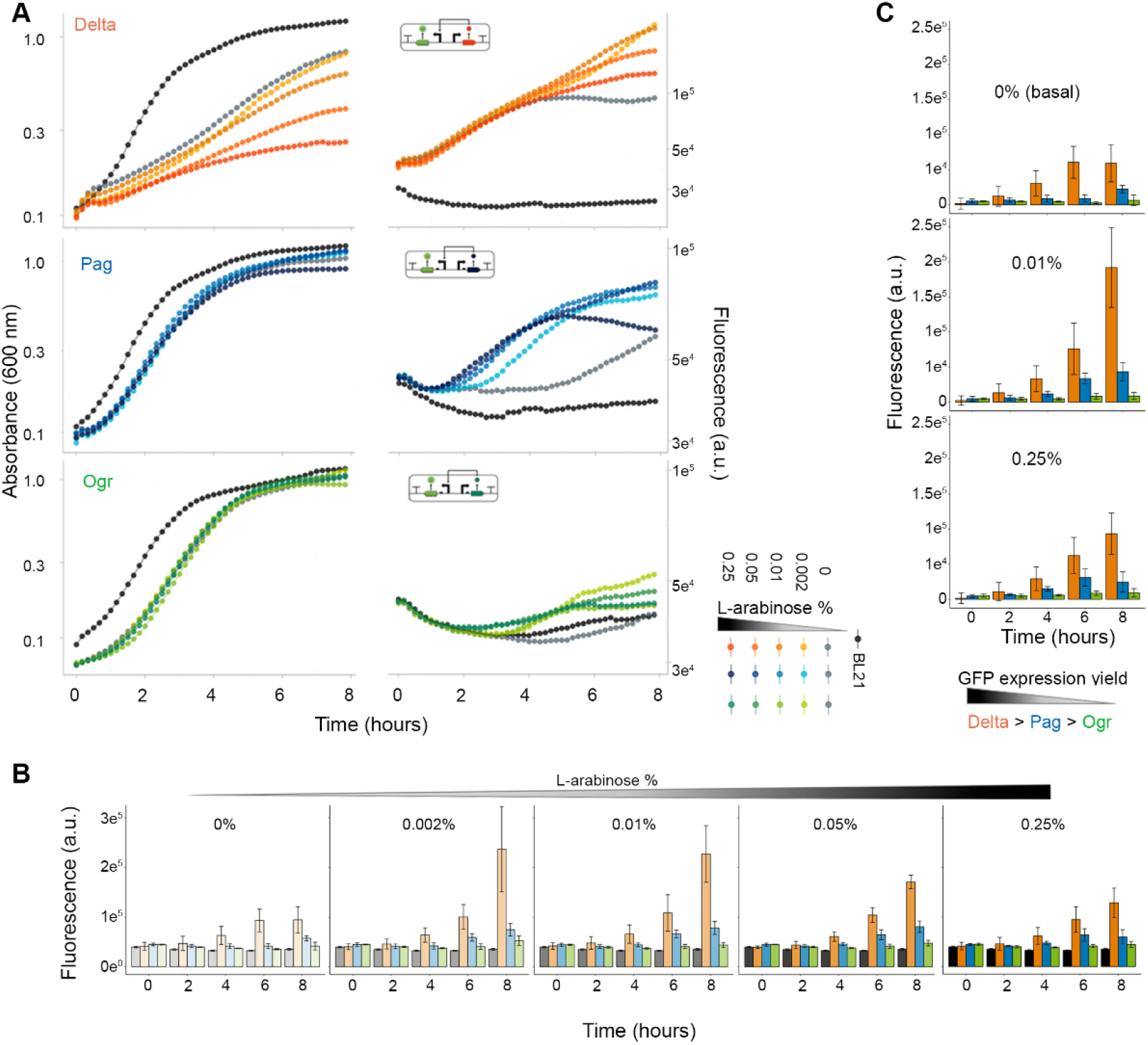
Characterization of the genetic circuits. **(A)** Growth curves (left) and time-dependent GFP expression monitored by fluorescence (right). The color code is designated on the schematic, with the strongest color representing the highest arabinose concentration. Non-transformed BL21(DE) cells are depicted in black. Data are means (n = 3 independent biological replicates). **(B)** GFP fluorescence shows dose-dependence on the arabinose concentration. Data are means ± SD (n = 3 independent biological replicates). **(C)** Normalized GFP expression to the value at the time point of arabinose addition (i.e. basal expression value without inducer). Data are means ± SD (n = 3 independent biological replicates).

The multilayer circuits were expressed and characterized in *E. coli* BL21(DE3) and assessed by the following criteria: (i) the burden they impose on the host by evaluating cell growth (Figure S2A), (ii) the output fluorescence signal from the GFP expression cascade (Figure S2B), and (iii) the strength of the GFP yield (Figure 2A). We monitored the time-dependent expression of GFP over 8 h, recording also the cell growth (Figure 2AB) and determined the dose dependence of the fluorescence yield on the arabinose (P_BAD_ promoter activator) (Figure 2B). Delta TF reached the highest GFP yield, however, it was the most detrimental to the cells growth, followed by Pag and Ogr (Figure 2A,B). It is important to note that all constructs exhibited a basal fluorescence even without an inducer (Figure 2BC), which is addressed in following sections. The most suitable concentrations of the arabinose inducer were between 0.01% - 0.25% (Figure 2C), that were further used for the sRNAs evaluation. Additionally, we evaluated the dose dependence of the expression of the multilayer circuits on the inducer by analyzing the behavior of the medium-strength circuit (Pag TF) with FACS (Figure S2C). The dose of the inducer controlled the first layer directly affected the yield of the fluorescence reporter and corroborated the concentration range (0.01% - 0.25%) which we previously selected by bulk fluorescence measurements (Figure 2A-C). We established 0.25% and 0.05% as high concentration, 0.01% as a medium, and 0.002% as a low dose of arabinose to induce the multilayer circuits.

### Design of synthetic sRNAs to modulating genetic circuits

Next, we designed synthetic sRNAs to inhibit TF, and consequently to decrease GFP expression (Table S1). The modulatory sRNAs were designed to target the mRNA of each of the three TFs (*Delta, Pag, Ogr*) with different repression strength (high - H, medium - M, low - L, control - C) (Figure 1A), which co-expressed with their cognate circuit ultimately resulted in four different circuits for each TF. The sRNAs were expressed under the control of the T7 promoter on a separate plasmid (Table S2) which was co-transformed in equimolar ratios with the corresponding plasmid expressing the multilayer circuit construct (Figure 1B). A dysfunctional sRNA (sRNA C) with a randomized scrambled sequence, that does not pair to the target was used as a negative control.

We next computed the duplex formation of the TF mRNA with the corresponding set of sRNAs (H, M, L, C). The predicted minimum free energy (MFE) values for all TF-sRNA complexes along with the predicted helical geometry of binding (Figure S3) indicated that as the sRNA modulating level decreased, the affinity of the sRNAs to the mRNA decreased and consequently, the duplexes were less stable.

### Assembling and debugging the multilayer genetic circuits

The multilayer circuits were subjected to three iterations of the classic ‘design-build-test’ cycle, which are referred to as generation 1, 2, 3 (G1, G2, G3) plasmids (Figure S4A, Table S2). The G1 generation was expressed from a pGGA plasmid. However, as determined by FACS the co-expression of G1 and plasmids bearing sRNA resulted in a noisy expression that was difficult to track (Figure S4B). To investigate whether the noise was generated from the sRNA or the genetic circuit plasmid, we designed a synthetic sRNA (high repression) to target an intrinsic TF (OmpR) of *E. coli*, with a binding affinity of -36.70 kcal/mol, and co-expressed it with wtGFP under the control of the *ompR* promoter. While we detected clear signal, we did not observe any suppression by this sRNA, because the fluorescence signals from wtGFP overlapped with that of the *ompR*-controlled GFP plus sRNA (Figure S4B).

Next, we performed a screening of the TF mRNAs for hairpins within their 5’UTRs that could prevent the synthetic sRNAs biding. We did not observe such structure that would alter the accessibility (Figure S5A). According to the predictions, a secondary structure formation for *Delta* TF mRNA is likely, however, the equilibrium probability of it was fairly low to persist. Using RT-PCR we detected the amplicon reporting on TF mRNA for Delta and Pag only after induction (Figure S5B). For Ogr TF, the same size amplicon was observed also in the uninduced sample, however, another primer pair yielded the same result (Figure S5B, C). The amplicon was most likely due to primers annealing, at least in part, to other places in the *E. coli* genome (Figure S5D).

By northern dot blot assay using tailored fluorescent probes (Table S3), we verified the expression of the sRNAs (H) (Figure S5E). After detecting an expression and discarding a possibility of failure in sRNA production, we reasoned that incompatibility of both plasmids could be an explanation for the noisy results (Figure S4B). Bacterial plasmids which share replication control or origin of replication (*ori*) are incompatible for tandem expression (Thomas & Smith, 1987; Novick & Hoppensteadt, 1978). Both G1 plasmids and sRNA plasmids were high copy plasmids with a pUC19-derived pMB1 *ori*. We next exchanged the plasmid backbone to a low-mid copy 15A *ori* plasmid (pSB3C5), producing G2 plasmids which however were still leaky, i.e. presenting high basal expression without induction (Figure S4C left), although there is an evidence that P_BAD_ promoter is tight (i.e. leaky transcription occurs at very low levels) (Fricke *et al*, 2020), and addition of *AraC* repressor should confer tight regulation (Guzman *et al*, 1995). Considering this, we created the G3 plasmids (Table S2). The GFP expression from both G2 and G3 plasmids exhibited dose-dependence on the inducer (arabinose) concentration (Figure S4C), yet the basal activity was still detectable (Figure 2C), corroborating earlier observations that the P_F_ promoter presents activity even without its activator (Julien & Calendar, 1996). Thus, in the characterization of the circuits, we subtracted the basal signal detected prior to induction.

### sRNAs robustly target TF expression to modularly control multilayer gene circuits

We evaluated the repression effect of the engineered sRNAs through different sets of experiments with varying conditions, thereby addressing the following questions: *i*) Are sRNAs functional or able to confer repression?, *ii*) Is the repression dose-dependent?, *iii*) Does the repression mirror the predicted repression strength?, *iv*) Is the repression effectivity dependent on the time of induction ? *v)* Does the predicted repression function holds across time?

First, we confirmed that the synthetic sRNAs are functional and capable of repressing GFP expression. Using the leakiest circuits (G2) at a basal expression (i.e. without arabinose induction), and triggering the sRNA expression at a maximum inducer concentration (2 mM IPTG), we observed a marked suppression of fluorescence as monitored by FACS (Figure S4D). Despite of the leakiness of the G2 circuits, we consistently observed the repression of all 3 TFs controlling the second layer of the circuits. sRNAs consistently showed a repression activity for both G2 and G3 circuits (Figure S4E), suggesting that the synthetic sRNAs are functional.

We further assessed the sRNAs activity with FACS, by analyzing the co-expression of the medium yield circuit (Pag TF) at a medium inducer concentration (0.05% arabinose) and with varying dosages of high repression (H) sRNA (Figure S2D, E). We observed a decrease in fluorescence, which is informative on efficient repression from sRNAs even at lowest concentrations of IPTG; a gradual decrease of the fluorescence signal as the IPTG concentration increased (Figure S2D). In turn, we did not observe a change in fluorescence when inducing the dysfunctional scrambled sRNA (Figure S2E), implying that the observed fluorescence decay was solely triggered by sRNA-dependent repression. Together these results suggest that synthetic sRNAs are functional and efficiently repress the TF genes in a manner dependent on the dose of the inducer (i.e. here IPTG).

Next, we assessed the dose-dependence of the IPTG inducer (in the range 0-2 mM) on the sRNA activity. This was evaluated with time-course fluorescence measurements in parallel to growth curves recording for all the circuits in tandem with the different sRNA plasmids (H, M, L, C). We used medium (0.01%, Figure S6) and high (0.25%, Figure S7) concentrations of arabinose (inducing the first layer), and we varied the IPTG concentration (0, 0.1, 0.25, 0.25, 1, 2 mM) to induce sRNA which was added at two different time points (e.g. 2h and 4h). Across all conditions tested, we detected a reduced growth of the cells carrying the multilayer circuits compared to the control (untransformed BL21(DE)). Thereby, the Delta-TF circuits were the most detrimental to the cells even without sRNA induction (e.g. 0 mM IPTG), suggesting that the synthetic circuits impose burden to the host. The production of sRNAs (0.1-2 mM IPTG) does not seem to additionally alter cell growth (Figure S6) even at high inducer concentrations (Figure S7). Overall, Delta TF, and the circuit with the highest GFP yield (Figure 2C), was the most difficult to control (Figure S6 and S7) which was independent of the level and time of induction. The repression effect was better detected when inducing the TF-GFP cascade at a medium (0.01% arabinose) level (Figure S6) and at early time point (2h) (Figures S6, S7).

We next addressed whether the predicted repression levels were achieved experimentally by measuring bulk fluorescence from cells (Figure 1A). Multilayered circuits and sRNAs were induced simultaneously with 0.002% arabinose and 2 mM IPTG, respectively. Fluorescence and cell growth were recorded 2h and 4h post induction (Figure 3A). Overall for all circuits, measured repression across distinct times post induction mirrored the predicted sRNA repression levels (i.e. H, M, L), except for Pag sRNA L at 4 h which yielded much lower fluorescence (or higher repression level) than sRNA M (Figure 3B). In all cases, sRNA H achieved the highest repression for all three circuits (up to 95%). It should be noted that we observed a modest repression activity (up to 12%) when using control sRNA.

**Figure 3.**
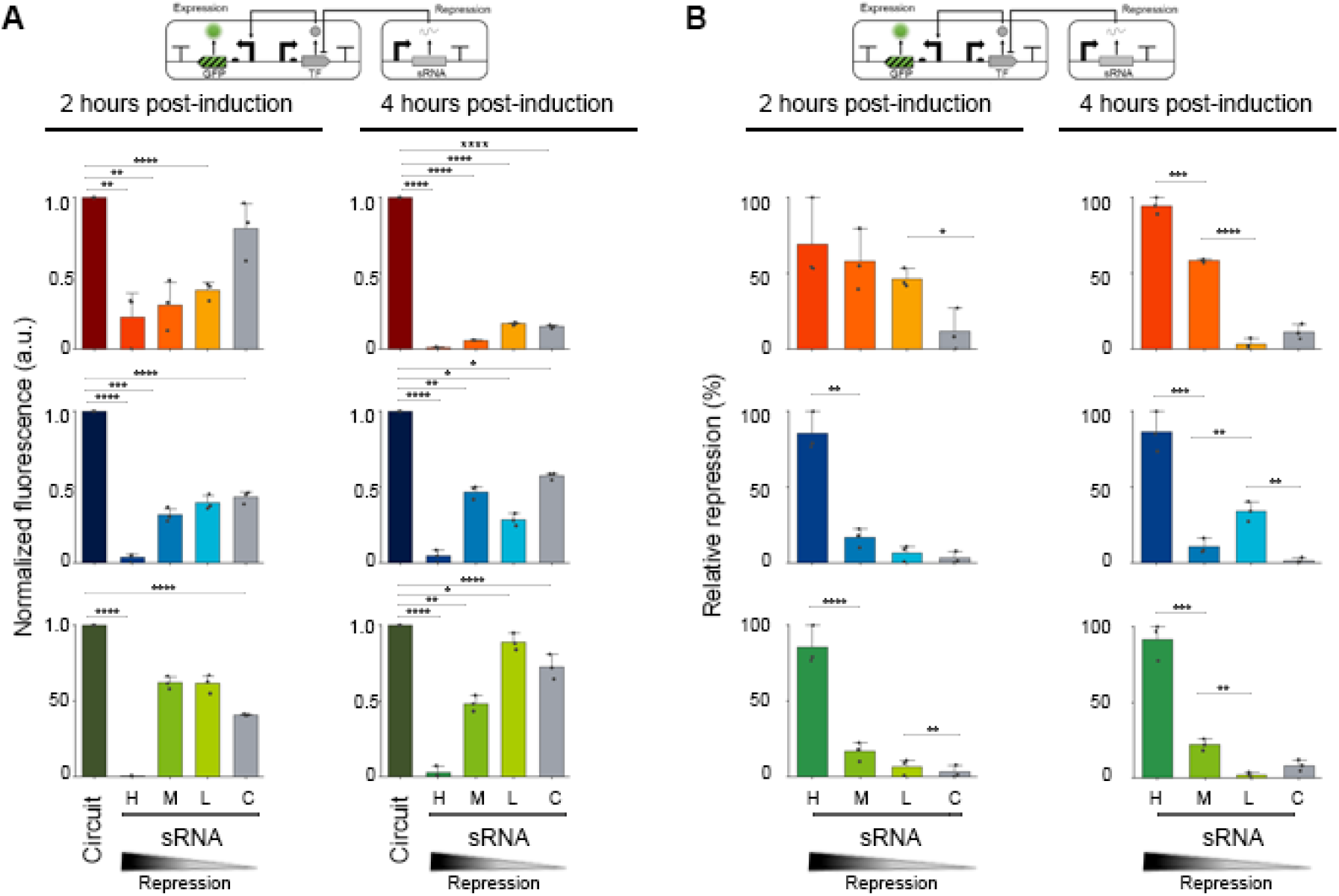
sRNAs efficiently modulate the expression level. **(A)** Change of bulk fluorescence intensity normalized to the circuit expression at 2 h and 4 h post induction. **(B)** Percentage of sRNA-mediated repression relative to the expression without sRNA. Delta – orange; Pag – blue; Ogr – green. Data are means ± SD (n = 3 independent biological replicates). *p < 1.0, **p < 0.05, ***p < 0.001, ****p < 0.0001, two-tailed t-test.

To assess the stability and efficiency of sRNA-mediated gene repression, we next monitored the GFP fluorescence decay after simultaneous induction of the multilayered genetic circuits and sRNA for 15 min (Figure 4). The time-course measurements were fitted to a two-phase exponential decay function with a time offset (Figure 4A), suggesting that two types, fast and slow exponential decay (likely after and before production of sRNAs), contribute to the signal. However, the rates of the slow and fast phase were similar (i.e. for Delta 0.039 and 0.037, for Pag 0.011 and 0.055, and for Ogr both 0.008 min^-1^, respectively). The data points before washing could not be fitted into one phase exponential decay as opposed to the data after induction, which was fitted remarkably well to a single exponential decay function (Figure 4B, C). We conclude that the drop of fluorescence resulted from the sRNAs-mediated TF repression on *GFP* mRNA, rather on the GFP protein turnover; the mature GFP has a long half-life (∼26 h), and by removing the inducers, no *de novo* protein is synthesized. The sRNAs repression efficacy was the highest for the Ogr circuit as suggested by the faster decay rates, followed by Delta and Pag sRNAs. Overall, the decay rates from the sRNA induced (or TF repressed) samples were consistently faster than the decay rates of circuits alone. The fluorescence half-life of sRNA-repressed circuits was lower compared to induced circuits even after removing the inducer (Figure 4D). These results support the notion that the fluorescence decay in circuits is effectively mediated by the synthetic sRNAs.

**Figure 4.**
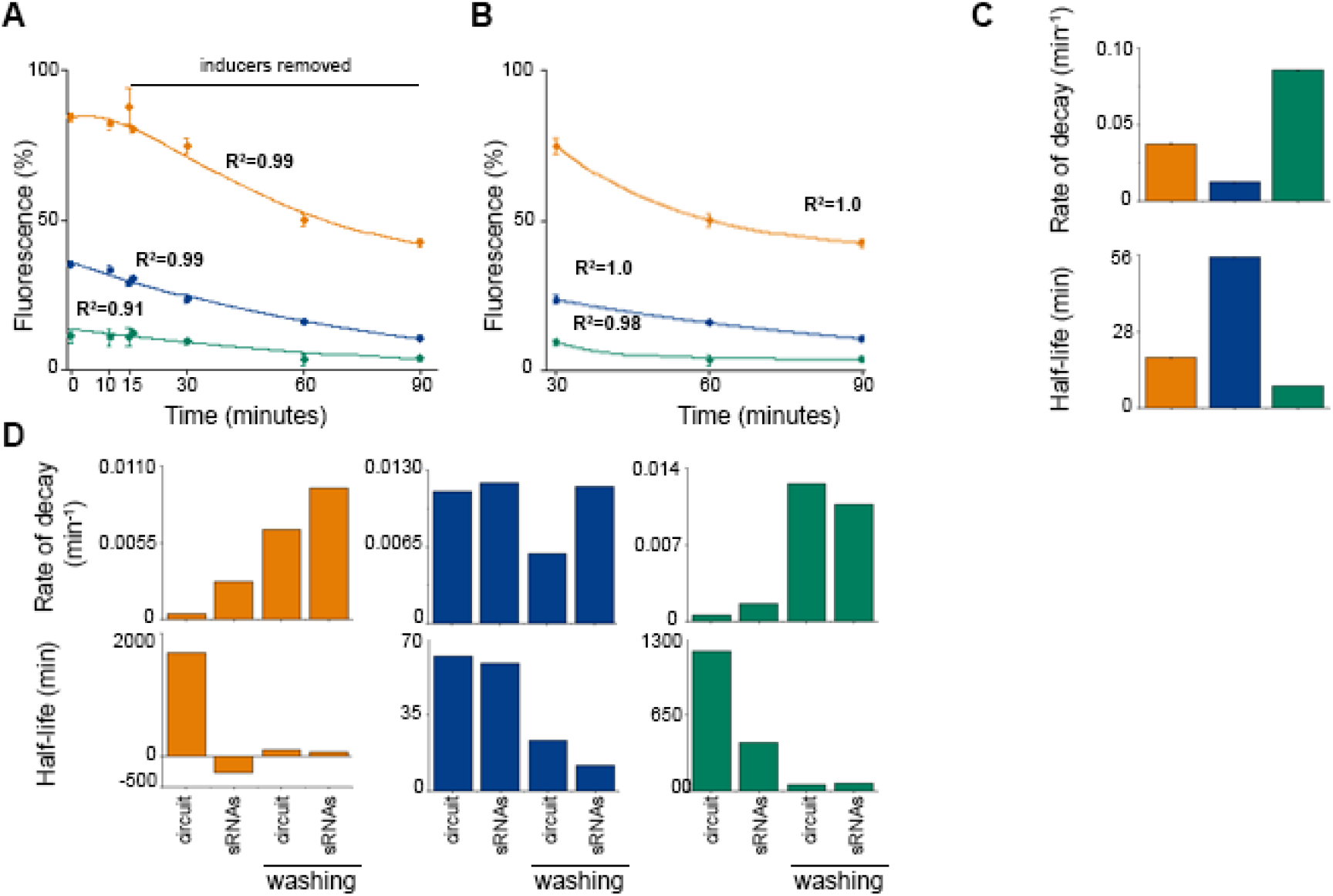
sRNA efficiently mediated the GFP fluorescence decay. **(A)** Time course of normalized GFP fluorescence (min-max normalized) by tandem expression of multilayered circuits and sRNA H, at medium (0.01%) and high inducer concentrations (2 mM), respectively; inducers were removed from the media at 15 min. Delta – orange; Pag-blue; Ogr-green. Curves were fitted to a two-phase exponential decay function. Data are means ± SD (n = 3 independent biological replicates). **(B)** Fluorescence measurements after depletion of the inducers (T=15 to T=90 min) fitted to a first order exponential decay function. Data are means ± SD (n = 3 independent biological replicates). **(C)** Decay rate and half-life of the fluorescence intensity derived from the fits to a first order exponential decay function (panel B). **(D)** Decay rate and half-life of fluorescence intensity before and after washing the inducers from the media, compared to induced multilayered circuit with and without sRNAs-expressing circuit.

## Discussion

Here, we investigated an sRNA-dependent modulation of multilayer genetic circuits as a means to post-transcriptionally control circuit’s output. De novo designed sRNAs target the effector of the second layer (i.e. a TF that controls the output of the circuit) and modulate its expression with different repression levels (H, M, L), thus allowing for a dynamic and precise fine tuning of the gene expression cascade. We addressed several important issues: *i*) Synthetic sRNAs are functional across multiple operational conditions (e.g. time, concentration of the inducer) and target different TFs, establishing different levels of repression; *ii*) sRNAs-based approach allows for modulating the off states dynamically, a state that is usually difficult to control; and *iii*) Experimental results robustly mirror the predicted values of repression with clear modular levels of repression from high to low (i.e. H, M, L). Our rationale is that sRNAs handle the repression of GFP upstream of its translation, and do not affect the concentration of already produced mature GFP. Mature GFP is remarkably stable and resistant to degradation (Kain, 1999), this altogether contributes to the remaining fluorescence.

In principle, our approach can be adapted to predictably modulate different genetic circuits by either targeting the effectors (e.g. such as TFs) or the circuit outputs. This notion is corroborated by the mathematical model describing the kinetics and reaction network of synthetic sRNA-mediated multilayer genetic circuit regulation (Figure S8, Table S5 and S6). Time-course simulations follow a similar trend as experimental results (Figure 4A) when our model was fitted with the decay rates for each TF (Figure S8C), suggesting the model could be further adapted to generically predict the regulation of genetic circuits sRNA-mediated regulation. It should be noted that not all mRNAs might be susceptible to the sRNA targeting. Previous studies have explored the tunability of genetic circuits at both transcriptional and translational level; examples include recombinant systems based on inducible promoters coupled with riboswitches (Morra *et al*, 2016) or regulatory motifs based on toehold switches paired to sRNAs (Bartoli *et al*, 2020). Other approaches for regulation of genetic circuits have paired artificial sRNAs with other techniques, such as CRISPRi (Cao *et al*, 2017), or riboswitches (Bartoli *et al*, 2020) to coordinate and control synergistically transcriptional and translational programs. In this study, we particularly explored the modulation of multilayer circuits by synthetic sRNAs targeting effector molecules (i.e. TFs), that in turn regulate the expression of the corresponding reporter molecule(s) (here GFP). By exploiting sRNAs sequence and binding affinities, we pioneer a new type of sRNA-mediated regulation of TF expression to control both modularly and dynamically multilayer genetic circuits. Previous strategies for obtaining different repression efficacy from synthetic sRNAs included variations in threir abundance through controlling expression using promoters with different strengths (Noh *et al*, 2019; Sung *et al*, 2016; Na *et al*, 2013). The construction of robust synthetic gene circuits, coupled with reliable sRNA-based approaches for their regulation, provides a framework for development of novel applications in the personalized medicine, RNA-based diagnostics and therapeutics, and improve strains for expression (Pardee *et al*, 2016; Harimoto *et al*, 2022; Noh *et al*, 2019).

## Materials and Methods

### Synthetic sRNAs

Target sequences were designed using in-house written scripts in Python (Figure S1). Binding energies were calculated using *RNAcofold* from the ViennaRNA package (Lorenz *et al*, 2011). The sRNA sequence was screened for off-targets against *E. coli* BL21(DE3) (Tax-id:866768) and str. K-12 substr. MG1655 (NC_000913) genomes, using RNAPredator (Eggenhofer *et al*, 2011). The first ten (ranked by base pairing affinity) interactions were selected for post-processing, the sRNA was selected there were no potential interactions found. Thereafter, the sRNAs sequences (DNA) were further screened with the web tool NEBcutter V2.0 to probe for restriction enzyme sites that could interfere with the cloning procedure. Target sequences were concatenated with the MicC scaffold (5’-TTTCTGTTGGGCCATTGCATTGCCACTGATTTTCCAACATATAAAAAGACAAG CCCGAACAGTCGTCCGGGCTTTTTTT -3’).

### Thermodynamic and secondary structure analysis

Minimum Free Energy (MFE) of the secondary structure of the TF mRNA-sRNA duplexes, their probability of formation, and helical geometry were calculated using NUPACK (Zadeh *et al*, 2011); 2D structure schemes were produced, as well. Parameters for the simulations were as follows: RNA energy parameter from Serra and Turner (1995); temperature was set to 37° C matching the optimum for *E. coli* experimental growth temperature; values for Na^+^ = 1.0 M, and Mg^++^= 0.0 M (set to default).

### Synthesis of components and assembly

For each TF, a genetic circuit plasmid plus three sRNA plasmids, one for different levels of sRNA-mediated repression (sRNA_H, sRNA_M, sRNA_L; see Table S1) were assembled. Multilayer genetic circuit plasmids were constructed using PCR and Golden Gate assembly (GAA) using *Bsm*BI Type IIS Enzyme (New England Biolabs), and sRNA plasmids using *Bsa*I Type IIS Enzyme (New England Biolabs) and transformed into *E. coli* strain DH5α. All sRNA plasmids were assembled with *Bsa1_micC_Suf_f* primer and a reverse primer with the specific sRNA sequence. New customized plasmids for sRNA expression can be constructed by using *Bsa1_micC_Suf_f* primer and a new reverse primer containing *Bsa*I recognition site at 5′ end, followed by an overhang (GAAA), the sRNA sequence (use reverse complement), and a binding sequence (complementary to the T7 promoter and the first 20-25 nt of the plasmid backbone). This is possible since the first four nucleotides of the *MicC* scaffold were always used as the overhang (TTTC). Plasmids and oligonucleotides used in this study are listed in Table S1 and S4, respectively.

### Characterization of multilayer genetic circuits and synthetic sRNAs

Circuit and sRNA plasmids were co-transformed into BL21(DE3) cells (i.e. Gx_TF + sRNA_TF_Lvl, where x=no.generation circuit, TF={Delta,Pag,Ogr}, Lvl={H,M,L,C}, see Table S2). Pre-cultures of transformed cells were grown overnight for 12−16 h at 37 °C 250 rpm with corresponding selection antibiotics. Main cultures were inoculated with the overnight cultures, and grown at 37° C in Luria−Bertani (LB) media (Roth, X968.4) supplemented with a 1:1000 ratio concentration of 100 mg/mL ampicillin (Sigma-Aldrich, A9393) and/or 34 mg/mL chloramphenicol (Sigma-Aldrich, C0378). Expression of TF circuits was induced with 0.002, 0.01, 0.05 or 0.25% (v/v) L-arabinose, respectively, and sRNAs were induced after 2 or 4 hours (post-circuit induction) with 0.1, 0.25, 0.25, 1, or 2 mM IPTG, respectively. Cell cultures were aliquoted for FACS (1 mL) and/or multiplate reader measurements (200 μL). For time-course assays, aliquots were grown and analyzed on at 37 °C with orbital shaking in a multimode microplate reader (Tecan Spark) using black 96-well plates with clear-flat bottom (Corning, Sigma-Aldrich CLS3603-48EA). Cell growth was measured at OD600 nm. GFP expression was measured in bulk (480 nm excitation/530 nm emission). Both measurements were taken every 10 minutes over a period of 8h, 12h or 14h (depending on the induction time) on a multimode microplate reader (Tecan Spark). To determine the basal fluorescence from the TF genetic circuits, both circuit and sRNA plasmids were co-transformed, however the expression of only TF circuits was induced (by adding only L-arabinose). Untransformed E. coli BL21(DE3) cells were grown to determine the background fluorescence, and for comparison as a control in the growth experiments. BL21(DE3) cells transformed with only circuit plasmids (i.e. Gx_TF, where x=no.generation circuit, TF={Delta,Pag,Ogr}) were also used as control. Min-Max normalization was applied to the data from all characterization experiments as follows: 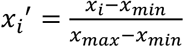, where *x*_*i*_ is the fluorescence value at a time point (*i*), *x*_*i*_′ is the normalized fluorescence value; *x*_*min*_, *x*_*max*_, correspond to the minimum and maximum fluorescence values on the complete data set. Calculations and plots were made by in-house R (version 4.2.1) written scripts.

### Modulating-repression of multilayer genetic circuits by sRNAs assays

The circuit plasmid and sRNA plasmids were co-transformed in E. coli BL21(DE3) cells and grown from overnight cultures at 37° C in LB media (Roth, X968.4) supplemented with antibiotics (1:1000 ratio concentration of 100 mg/mL ampicillin (Sigma-Aldrich, A9393) and/or 34 mg/mL chloramphenicol (Sigma-Aldrich, C0378)). When the cultures reached approximately an OD_600_ of 0.5, circuit expression was induced with a medium dosage (0.01% L-arabinose) and sRNAs with the maximum dosage (2 mM IPTG). Co-transformed cells induced with only L-arabinose (only circuit-no sRNAs), transformed cells (only circuit plasmid) and blank BL21(DE), were used as control. Cells were cultivated, and aliquots were taken at 2h and 4h post induction for bulk fluorescence (200 μL) and FACS (1 mL) analysis. Bulk fluorescence measurements (480 nm excitation/530 nm emission and optical density 600 nm) were performed on a microplate reader (Tecan Spark) using a black 96-well plate with clear-flat bottom (Corning, Sigma-Aldrich CLS3603-48EA). Data was analyzed as follows: absolute change of intensity (*F*_*∆*_) from the repressed circuits with respect to unrepressed circuits was calculated from absolute fluorescence values (*F*_*abs*_) and were normalized to the unrepressed circuit fluorescence (i.e. taken as maximum fluorescence). Absolute fluorescence values from all samples (*F*_*sample*_) were calculated by taking into account the background fluorescence value from culture medium (*F*_*LB*_) as follows: 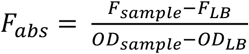. Absolute change of intensity (*F*_*∆*_) in fluorescence with respect to the circuit was calculated as follows: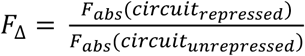. Relative repression percentage of sRNAs was calculated by expressing the relative change in fluorescence from unrepressed circuits with respect to repressed circuits in percentage: 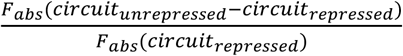. The statistical analysis of the data was performed with a 2-tailed t-test, calculated with in-house Python (version 3.9.10) and packages NumPy version 1.22.3, and SciPy version 1.8.0 scripts.

### Fluorescence activated cell sorting (FACS)

Aliquots from cells subjected to analysis were harvested, sedimented and washed with cold PBS (500 μL). After washing, cells were sedimented and resuspended in 1mL cold PBS. Samples were diluted (1:100; 900 μL PBS, 100 μL sample) and measured by a flow cytometer (FACSCalibur, Becton Dickinson) with CellQuest™ Pro Software Version 6.0. The measurements of GFP fluorescence were taken for ∼100,000 events with the following settings: FSC = E01, SSC = 400, FL1 = 736, in log scale and threshold FSC = 52. Histograms were plotted with FCSalyzer 0.9.22.

### Synthetic sRNAs decay assay

Circuit and sRNA plasmids were co-transformed and grown at 37° C in LB media (Roth, X968.4) supplemented with antibiotics (1:1000 ratio concentration of 100 mg/mL ampicillin (Sigma-Aldrich, A9393) and/or 34 mg/mL chloramphenicol (Sigma-Aldrich, C0378)). When OD_600_ of 0.5 was reached, the circuit expression was induced with a medium dose (0.01%) L-arabinose and sRNAs with the maximum dose IPTG (2 mM). After 15 mins, cells were harvested and washed twice with cold PBS (500 µL). Cells were resuspended in fresh LB (same volume) and grew further, aliquots (200 µL) were withdrawn at 30 min, 60 min- and 90 min. Optical density (600 nm) and GFP fluorescence were quantified in bulk (480 nm excitation/530 nm emission) with a microplate reader (Tecan Spark) using a black 96-well plate with clear-flat bottom (Corning, Sigma-Aldrich CLS3603-48EA). The full data set from the decay assay (i.e. before and after washing inducers, Figure 4A) was fitted to a two-phase exponential decay function with time offset of the form 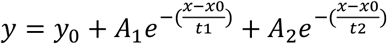 were *y*_0_ is the time offset, *t*_1_, *t*_2_ the time constants; *x*_0_ the peak center, and *A*_1_, *A*_2_ the amplitudes of the fitted curve. Fluorescence percentage data after washing the inducers (Figure 4B) was fitted to a one-phase exponential decay function with time constant of the form *y* = *y*_0_ + *Ae*^−(*x*/*t*)^ were *y*_0_ corresponds to the time offset, *t* is the time constant; *x* the peak center, and *A* the amplitude of the fitted curves. Exponential fittings were performed with OriginLab Pro 2019 (version 9.6.0.172). The rate of decay (*k* = 1/*t*_*x*_) and half-life (*T*_1/2_ = *t*_*x*_ *ln*(2)) were calculated as derived parameters of the fitting results (Figure 4C). Details on parameters values are specified in supplementary data files.

### RNA isolation and northern dot blot

A tailored protocol for RNA isolation was used and optimized to obtain a concentration of total RNA of approximately 300 ng/µl from 1:40 diluted samples (or 12,000 ng/µl in the undiluted samples). Single colonies of *E. coli* BL21(DE3) cells transformed with the sRNA plasmids were cultured in 1.3 L of LB media (Roth, X968.4) supplemented with 1:100 ratio of 100 mg/mL ampicillin (Sigma-Aldrich, A9393) at 37° C until OD_600_ = 0.5 was reached. Each culture was separated into 2 cultures (600 mL), one of them induced with 2 mM IPTG (sRNA production) and the other one used as control (no sRNA), therefore uninduced and induced samples handled under the same culturing conditions.

Cells were harvested (from 300 mL cell culture) at 2h and 4h post induction. Total RNA was isolated with TRIzol Reagent™ Solution (ambion, 15596018) and chloroform:isoamyl alcohol 24:1 (Sigma-Aldrich) accordingly to manufacturer’s instructions. To detect the production of sRNAs, Cy3 labeled oligonucleotides (Table S3) were used as probes for sRNAs, and 5S rRNA (as a control). Total RNA from untransformed BL21(DE) cells, which do not express sRNAs, was also subjected to northern dot blotting and used as negative control. Equal volumes (2 µL) of total RNA (∼10 µg) and denaturation buffer were incubated for 2 min at 95 °C. Denatured samples were directly spotted onto a nylon membrane (Amersham Hybond - N+ (0.45 µm)), crosslinked to membrane (exposed to UV light 254 nm, 120 mJ, twice), and incubated in Church’s buffer for 2h at 42 °C with rotation. Fluorescent probes (5 µL) were denatured at 90° C for 5 minutes, and added to the membranes. Membranes were probed with the corresponding DNA overnight at 28 °C in Church’s buffer. Fluorescence intensity of the hybridized probes was recorded on a Universal Hood III imager (Bio-Rad) with Image Lab™ software version 5.2.1(Bio-Rad) using the default blot protocol for Cy3 channel. Aliquot of each sample was kept for total counterstaining using methylene blue.

### RT-PCR

RT-PCR was used to detect TF transcripts and verify their expression. Cultures were induced at OD_600_ = 0.05 with 0.25% L-arabinose, then grown for 90 minutes. Selected samples were induced with 0.1% L-arabinose and incubated for 120 minutes at 37° C. Total RNA was isolated using TRIzol Reagent™ Solution (ambion, 15596018). Thereafter, a standard protocol for cDNA conversion by RevertAid Reverse Transcriptase (ThermoFisher Scientific, EP0442) was followed. The cDNA served as a template for PCR amplification using the corresponding primers (Table S4). Amplicons were verified by a 3% agarose gel electrophoresis stained with RedSafe™ (iNtRON, 21141).

### Mathematical model of sRNA-mediated genetic circuit regulation

Ordinary differential equations describing the kinetic model as follows:

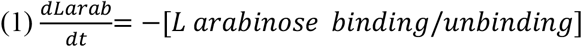

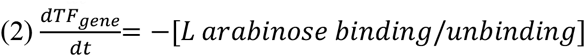

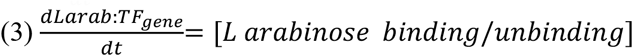

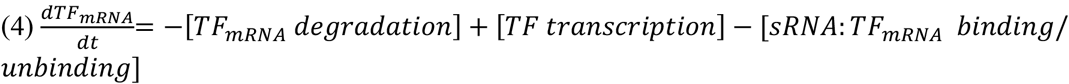

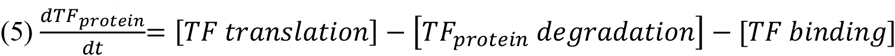

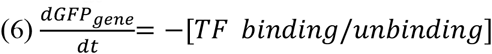

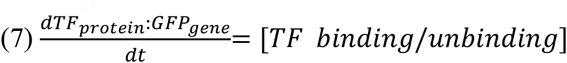

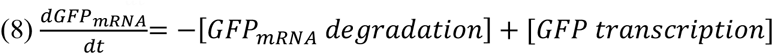

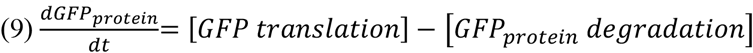

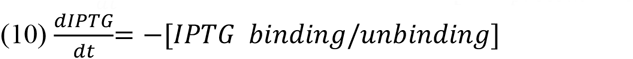

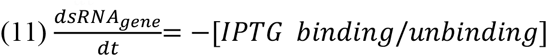

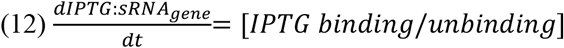

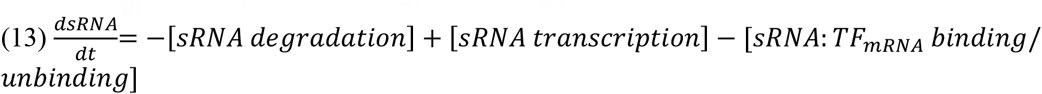

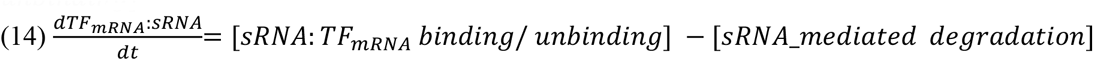

Equations 1-9 describe the first and second layers of the circuit, i.e. the production of a TF that in turn triggers the production of the circuit output (here GFP). Equations 10-14 model the sRNA production and their effect on the TF mRNA, and consequently in the GFP concentration. Repression by sRNAs on the TF mRNA was modeled as a receptor-ligand degradation reaction (reaction 15). The promoter activation from inducers (arabinose, IPTG) and TF were modeled as reversible binding reactions following receptor-ligand kinetics (reactions 1, 11 and 6 respectively). The secondary layer of the genetic circuits, when TF is produced and activates GFP, is described by reactions 1-10. The third layer, corresponding to the sRNA production and the decay of the TF mRNA mediated by the sRNAs is explained by reactions 11-15. Simulations of the mathematical model of the reaction network (Figure S8) were performed with SimBiology module from MATLAB (R2022a Update 4 version 9.12.0.2009381). Parameters and reaction rates (Table S6) were taken from the base model for gene expression and further modified to fit the reactions of the system (Table S5). Deterministic simulations were performed with the Suite of Nonlinear and Differential/Algebraic Equation Solver (sundials). To indicate the absence of sRNA production initial value of IPTG was set to 0. For the model to reflect our experimental findings, we set the parameter rate of the sRNA-mediated decay reaction (i.e. k_15_) to the maximum decay rate obtained from the fluorescence decay assay (i.e. Ogr; Figure 4C). To adjust different strength each TF exerts on the PF promoter, the GFP transcription rate (k_7_) was altered. Rates of sRNA decay obtained from experiments (Figure 4C) were plugged in by updating the value of the corresponding parameter (k_15_) expressed in sec^-1^.

## Acknowledgements

This work was supported by CONACYT-DAAD fellowship to A.K.V.S.

## Autor contributions

**Ana Velazquez:** Conceptualization; formal analysis; investigation; visualization; methodology; writing – original draft; writing – review and editing. **Bjarne Klopprogge**: investigation. **Karl-Heinz Zimmermann**: Conceptualization; supervision; writing – review and editing. **Zoya Ignatova:** Conceptualization; resources; supervision; writing – original draft; writing – review and editing.

## Conflict of interest

The authors declare no conflict of interest.

## Supplementary Information for

**Figure S1.**
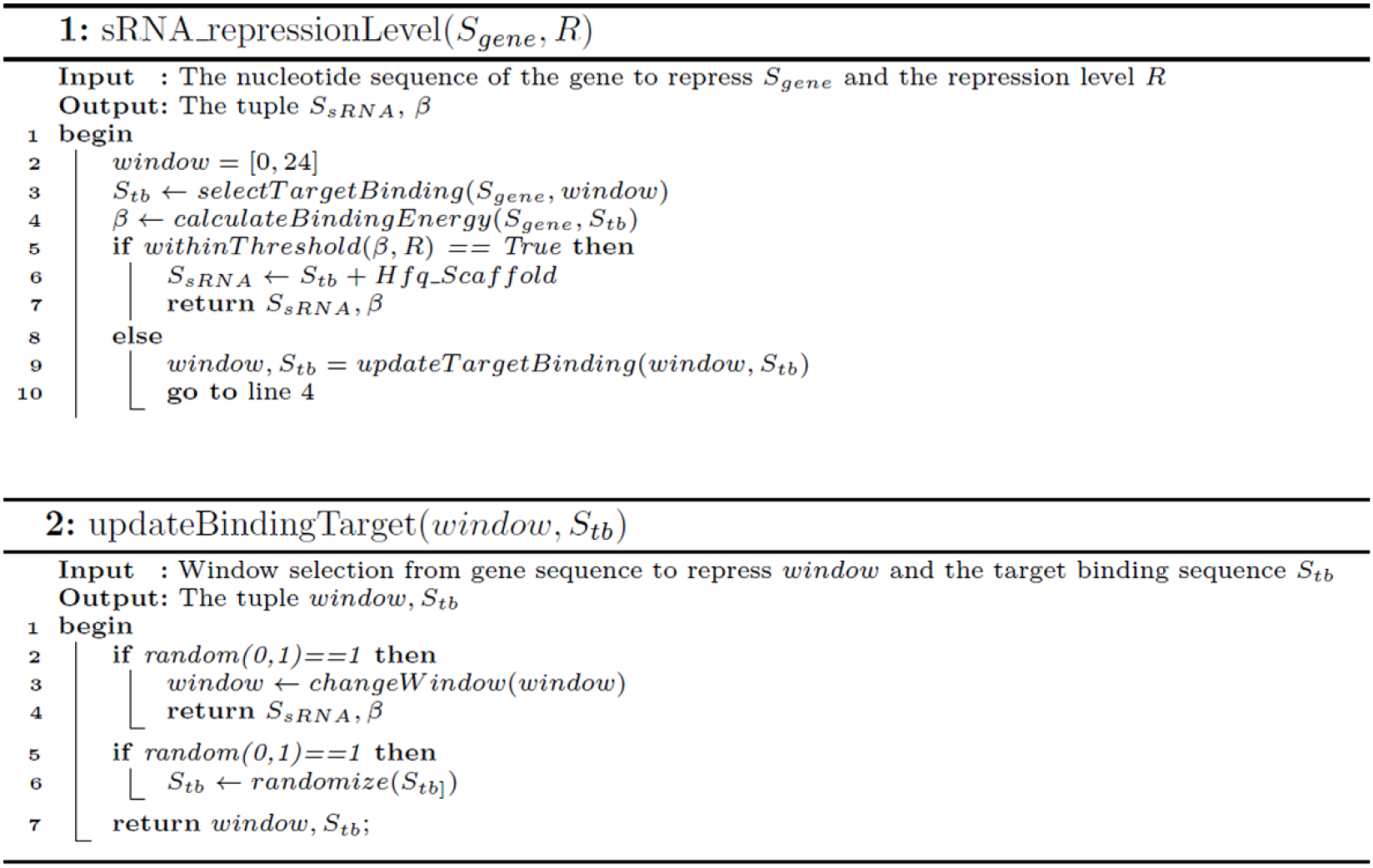
Algorithm to design synthetic sRNAs with a defined level of repression. *S*_*gene*_ is the sequence of the gene to repress; *S*_*sRNA*_ the tailored sRNA with the specified repression level; *β* the binding energy between the sRNA and the mRNA of the gene; *S*_*tb*_ the sequence of the target binding sequence of the sRNA (excluding the Hq scaffold); *R* = {*h, m, t*} the repression level of the sRNA h: high, m: medium, l: low; *Hfq_scaffold* the nucleotide sequence of the Hfq scaffold from Mic C (see Methods).

**Figure S2.**
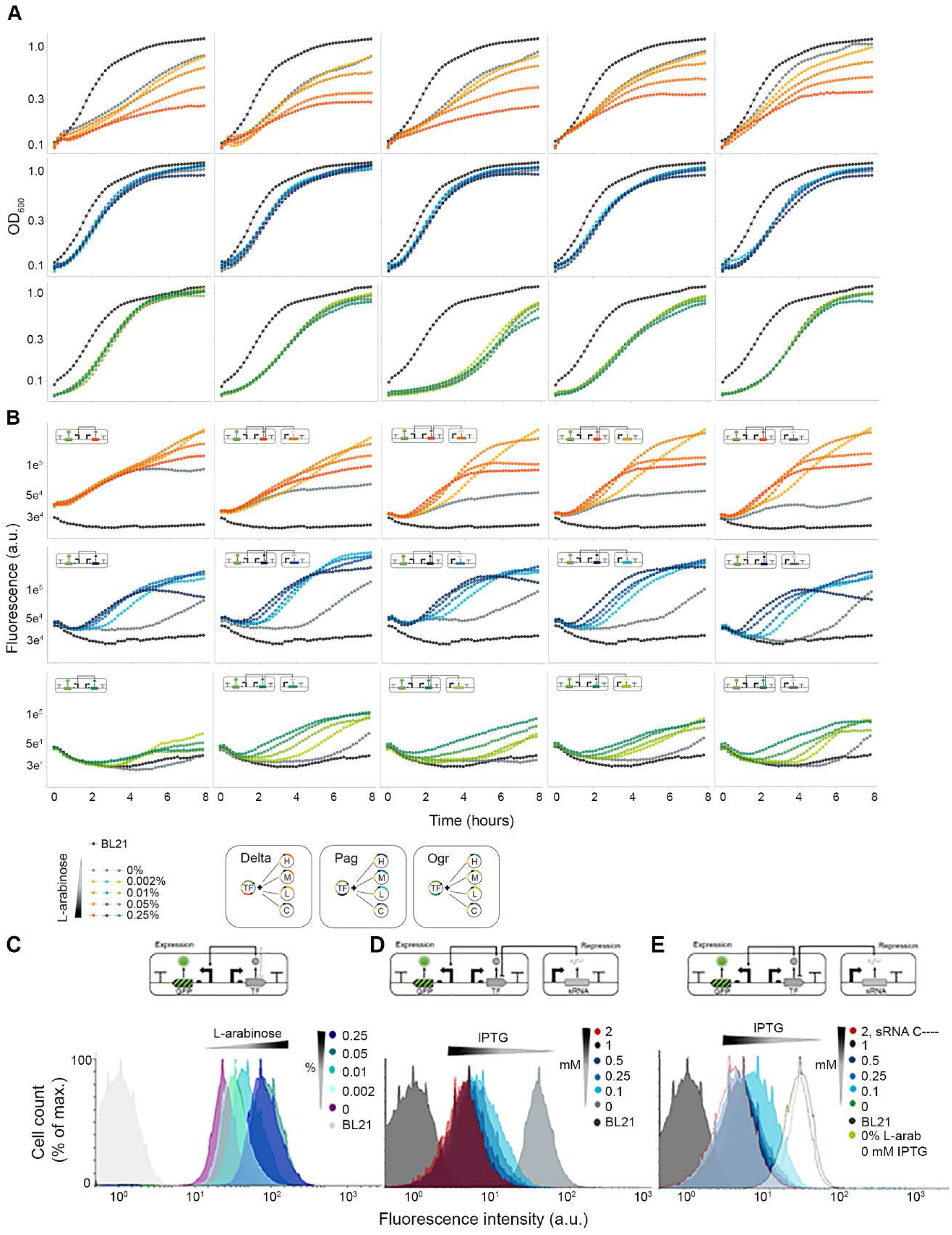
Output yield of circuits and sRNAs repression depend on inducers concentration. **(A, B)** Burden imposed by circuit plasmid (first column) and tandem expression of each circuit with plasmids expressing different sRNA levels (uninduced) analyzed by cell growth measured by optical density (**A**) and normalized mean fluorescence **(B)** from three biological replicates at different concentrations of arabinose. Delta-orange; Pag-blue; ogr-green. **(C-E)** Pag multilayer circuit (medium GFP yield) cascade performance at different concentration of arabinose with **(C)** induced at high level (0.05% arabinose) and varying level of sRNA H (high repression); **(D)** induced at high level (0.05% arabinose) and varying levels of sRNA H (high repression; **(E)** Leakiness tested with circuit without inducer. Green, sRNA repression; dashed, circuit and control sRNA repression.

**Figure S3.**
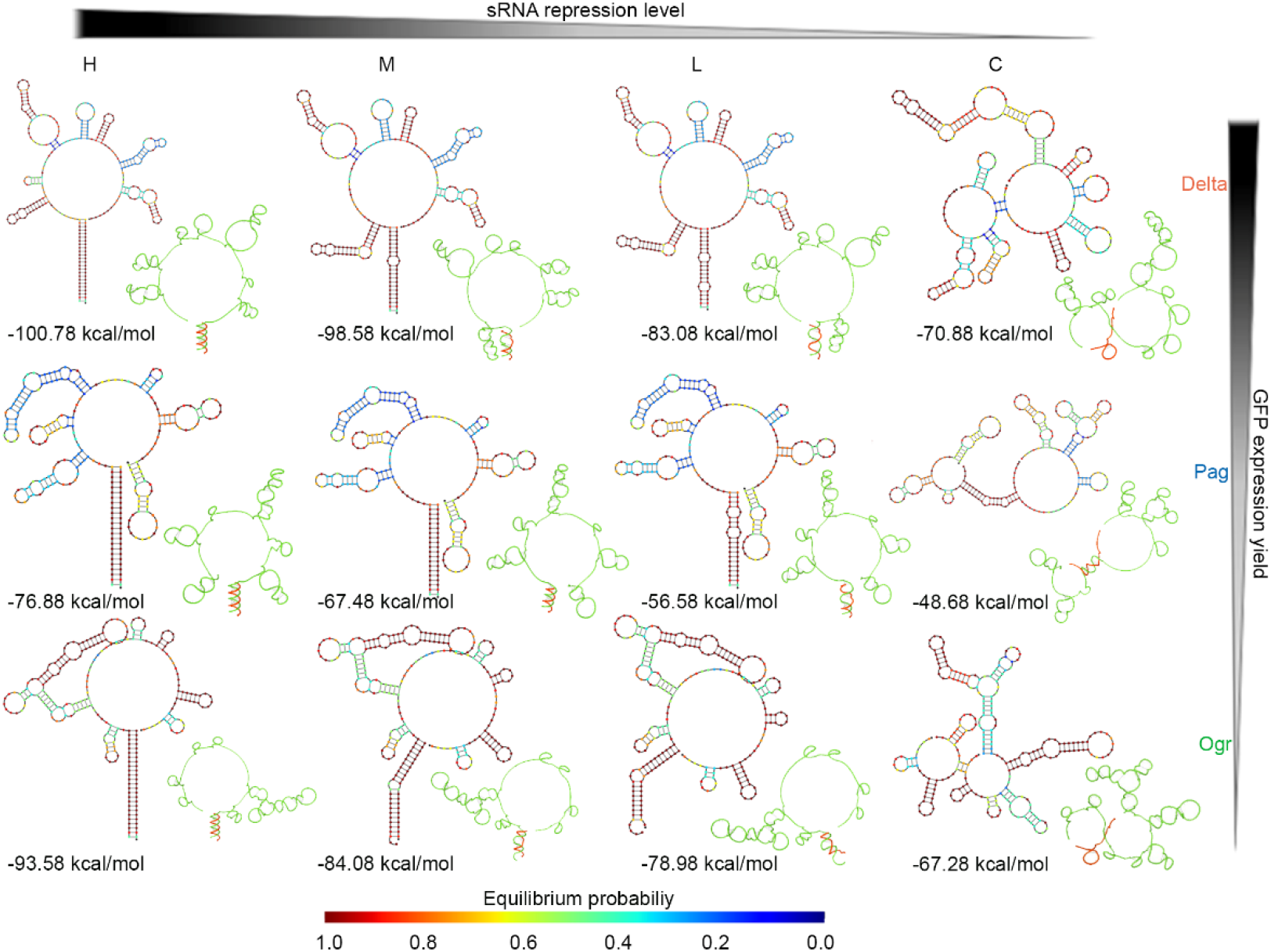
Thermodynamic analysis of sRNA-mRNA duplexes supports their different levels of repression. Minimum Free Energy (MFE) structures, ideal helical geometry (right), and free energy of the MFE (bottom) of *delta, pag, ogr* mRNAs paired with their cognate sRNAs with different translational regulation strength: H, high repression; M, medium repression; L, low repression; C, scrambled control. Predictions were performed with NUPACK. The color key indicates the probability of base pairing. The helical structure predictions depict the mRNA (green) and the sRNA (red) duplexes.

**Figure S4.**
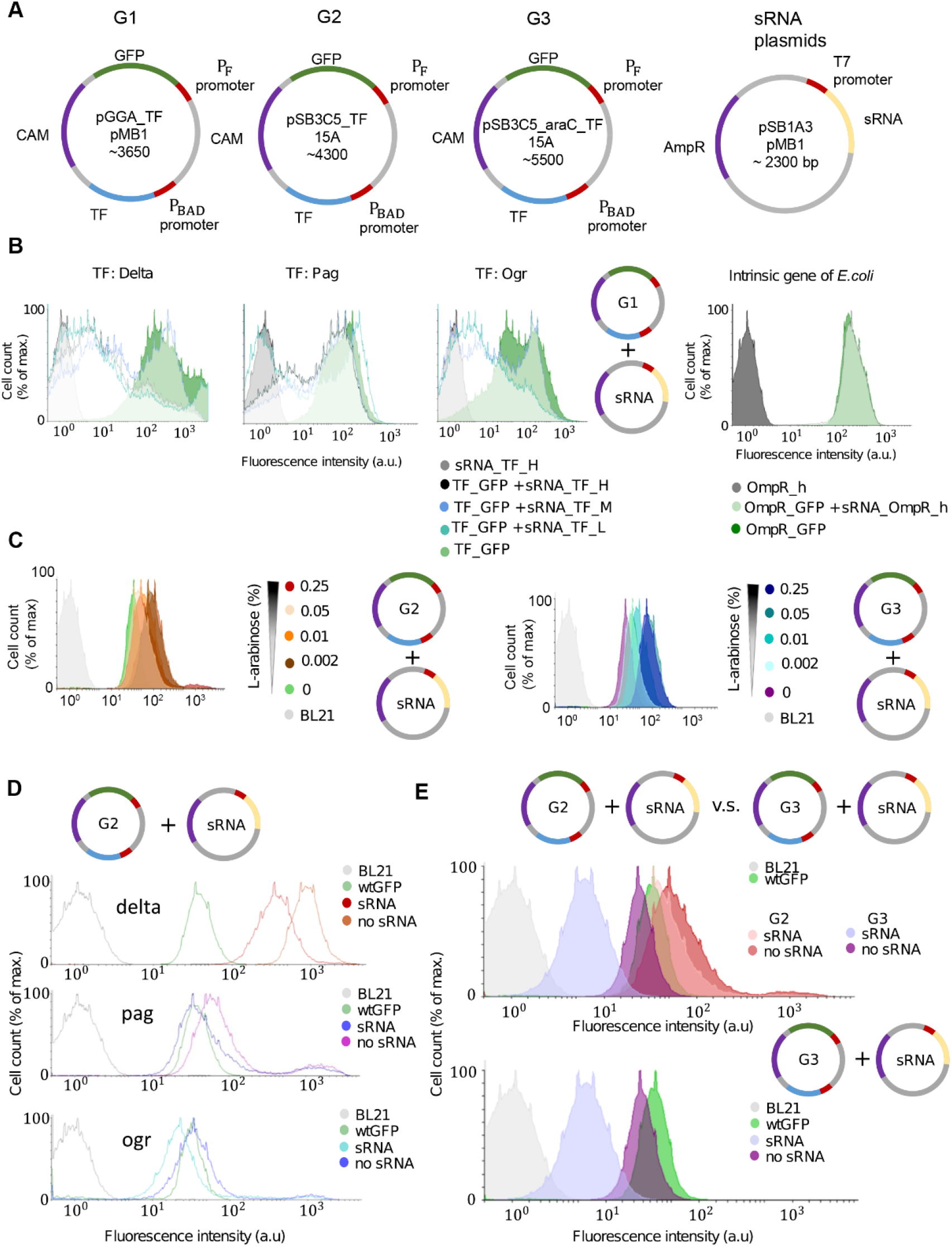
Performance of multilayered circuits evaluated by FACS. **(A)** Different generations (G1, G2 and G3) of multi-layered genetic circuit plasmids. **(B)** Tandem expression of genetic circuits G1 and sRNA plasmids and comparison of a genetic circuit induced by an intrinsic *E. coli* gene (*ompR*) and circuit regulation by sRNAs. **(C)** G2 circuits dosage compared to G3 circuits dosage dependence. **(D)** Tandem expression of all G2 circuits (without induction - leakiness) and synthetic sRNA (highest repression). **(E)** Comparison of performance of tandem expression between G2 and G3 circuits (without induction - leakiness) and sRNA (high repression).

**Figure S5.**
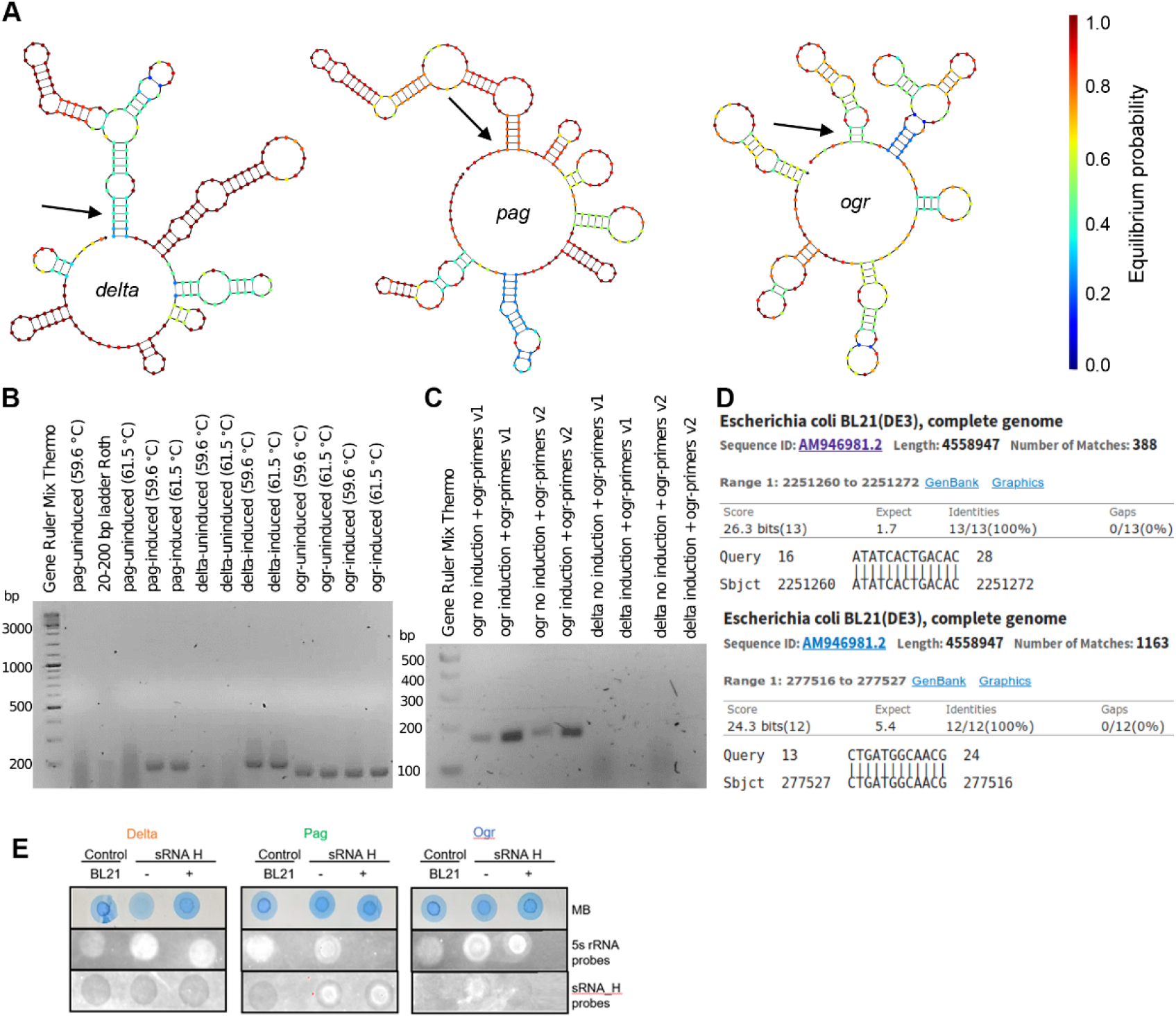
Debugging strategies for optimizing the co-expression of multilayer circuits and sRNAs. **(A)** Secondary structure prediction of TF mRNAs with 5’UTR pointed by black arrow. The color key indicates the probability of base pairing. **(B)** cDNA amplicons of TFs visualized on 3% agarose gel. All TFs display fragments of expected size. **(C)** Alternative primer pairs tested for *ogr* samples. Note that same amplicons were detected in the uninduced sample (ogr no induction) with both primer pairs. **(D)** BLAST alignment of the forward and reverse primers used to amplify *ogr* to the *E*.*coli* genome. **(E)** Assessment of sRNA high expression (sRNA_h) probed with specific probe by northern dot blots; MB, methylene blue staining of the total RNA used to probe the equal RNA amounts loaded onto the blot manifold. Probing of the 5S rRNA served as a control. Probe sequences are included in Table S3.

**Figure S6.**
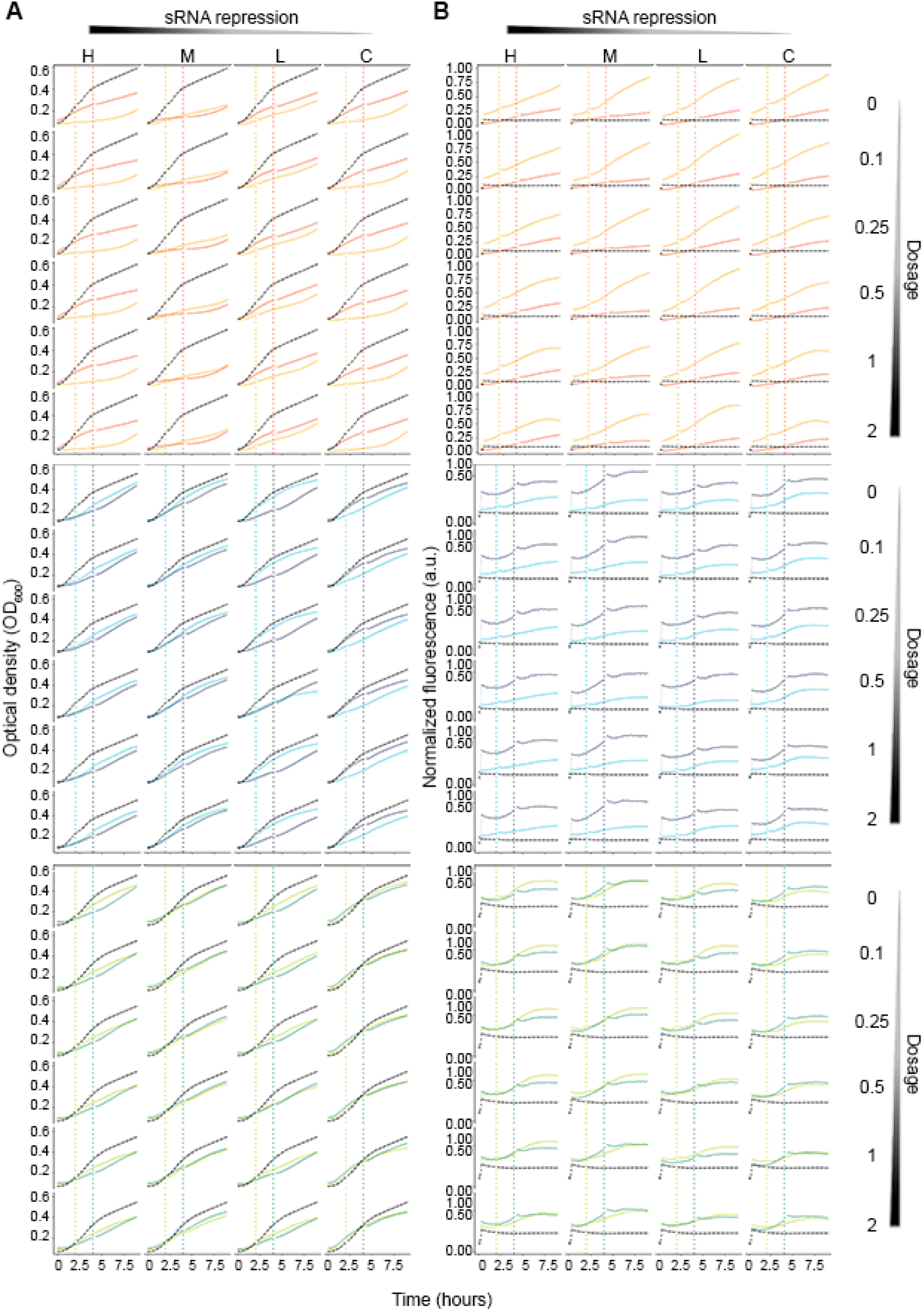
Real-time characterization of tandem expression of multilayer circuits and sRNAs. **(A, B)** Burden imposed by tandem expression of circuits and modulating sRNAs monitored by growth **(A)** or GFP fluorescence **(B)**. TF-GFP construct were induced at medium dose of arabinose (0.01%), and IPTG concentration varied. Comparison of the effect of all sRNAs targeting different TFs (Delta-orange, Pag-blue, Ogr-green) at early induction (2 h - light color) and late induction (4 h - dark color). Fluorescence values are normalized mean fluorescence from three biological replicates. Untransformed *E. coli* BL21 is shown in black.

**Figure S7.**
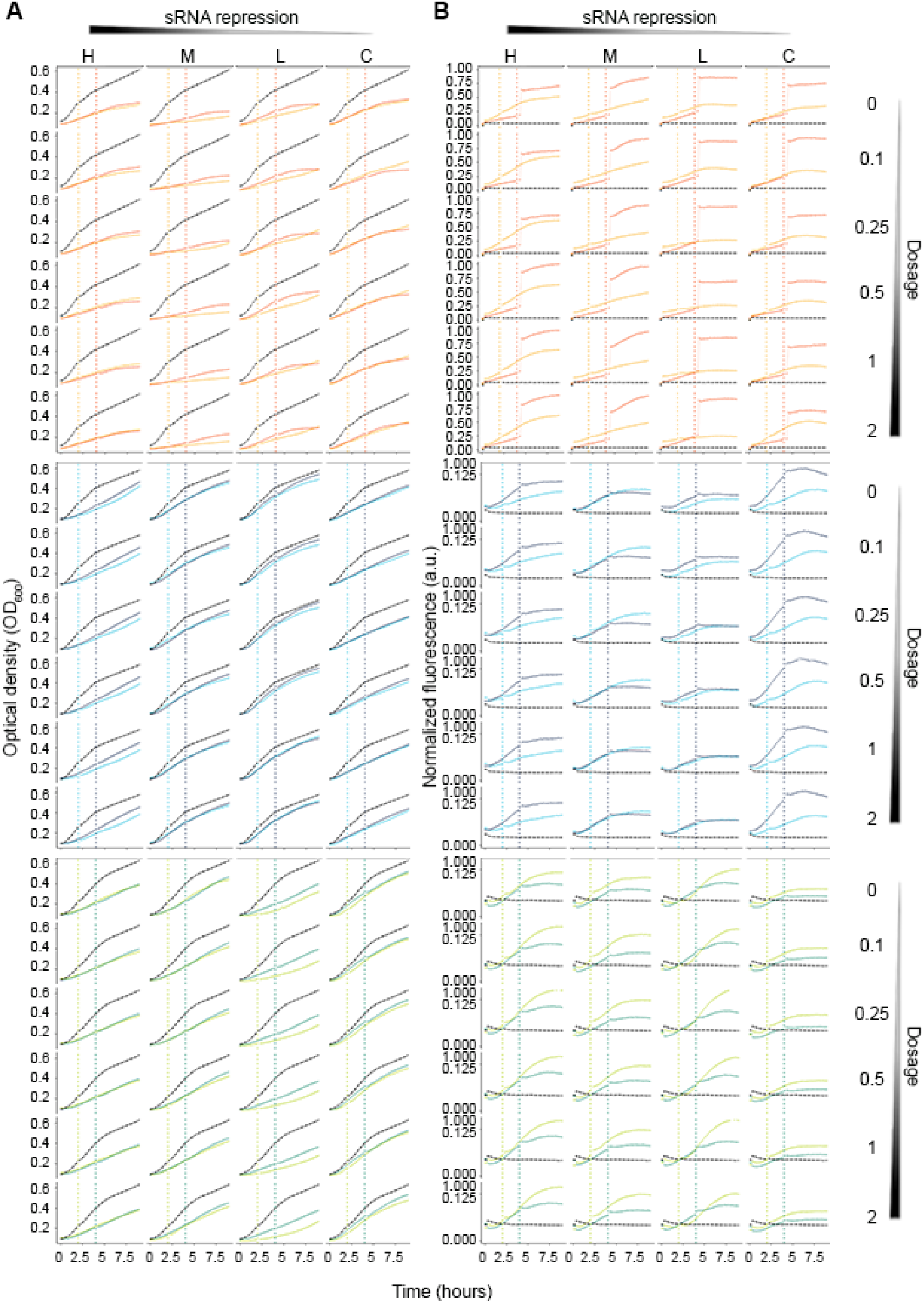
Real-time characterization of tandem expression of multilayer circuits and sRNAs. **(A, B)** Burden imposed by tandem expression of circuits and modulating sRNAs monitored by growth **(A)** or GFP fluorescence **(B)**. TF-GFP construct were induced at medium dose of arabinose (0.25%), and IPTG concentration varied. Comparison of the effect of all sRNAs targeting different TFs (Delta-orange, Pag-blue, Ogr-green) at early induction (2 h - light color) and late induction (4 h - dark color). Fluorescence values are normalized mean fluorescence from three biological replicates. Untransformed *E. coli* BL21 is shown in black.

**Figure S8.**
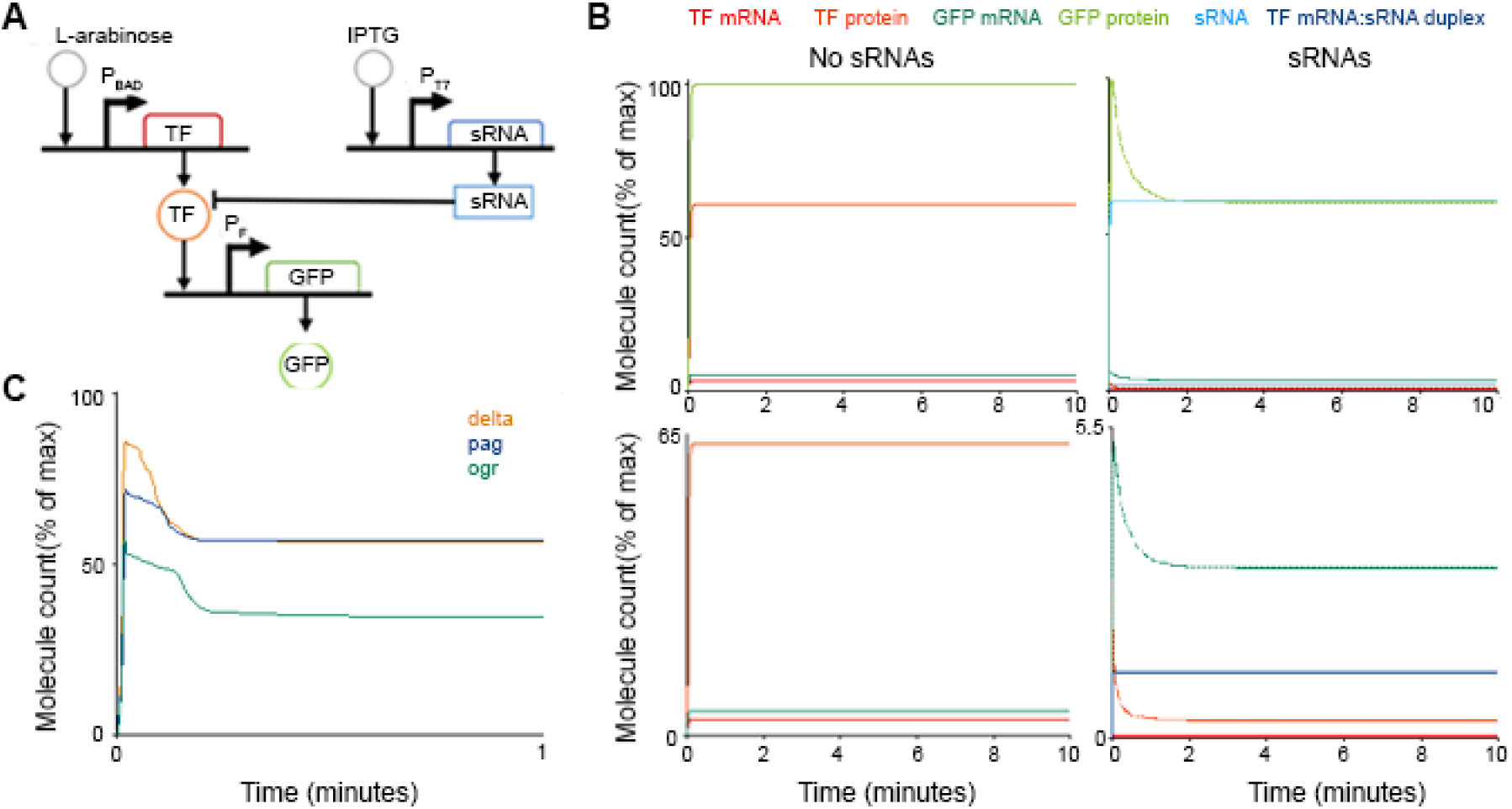
Modelling synthetic sRNA-mediated genetic circuit regulation. **(A)** Schematic of the reaction network used in the model. **(B)** Time-course simulations of the genetic circuit regulation mediated by synthetic sRNAs. Top plots comprise all species at full scale, bottom plots are zoom-in for species of interest. **(C)** Time-course simulation of the experimental results from the GFP fluorescence decay (Figure 6C). The following parameters were used: delta (k_7_=0.7 sec^-1^, k15=2.25 sec^-1^); Pag (k_7_=0.5 sec^-1^, k15=0.75 sec^-1^); Ogr (k_7_=0.4 sec^-1^, k_15_=5.11 sec^-1^). Deterministic simulations mimic the experimental findings, i.e. a substantial decrease in GFP expression compared to conditions with sRNAs not being produced (B). The drop of the TF protein and GFP mRNA indicates that GFP production is inhibited by the reduced TF concentration (B, bottom right panel).

**Table S1.**
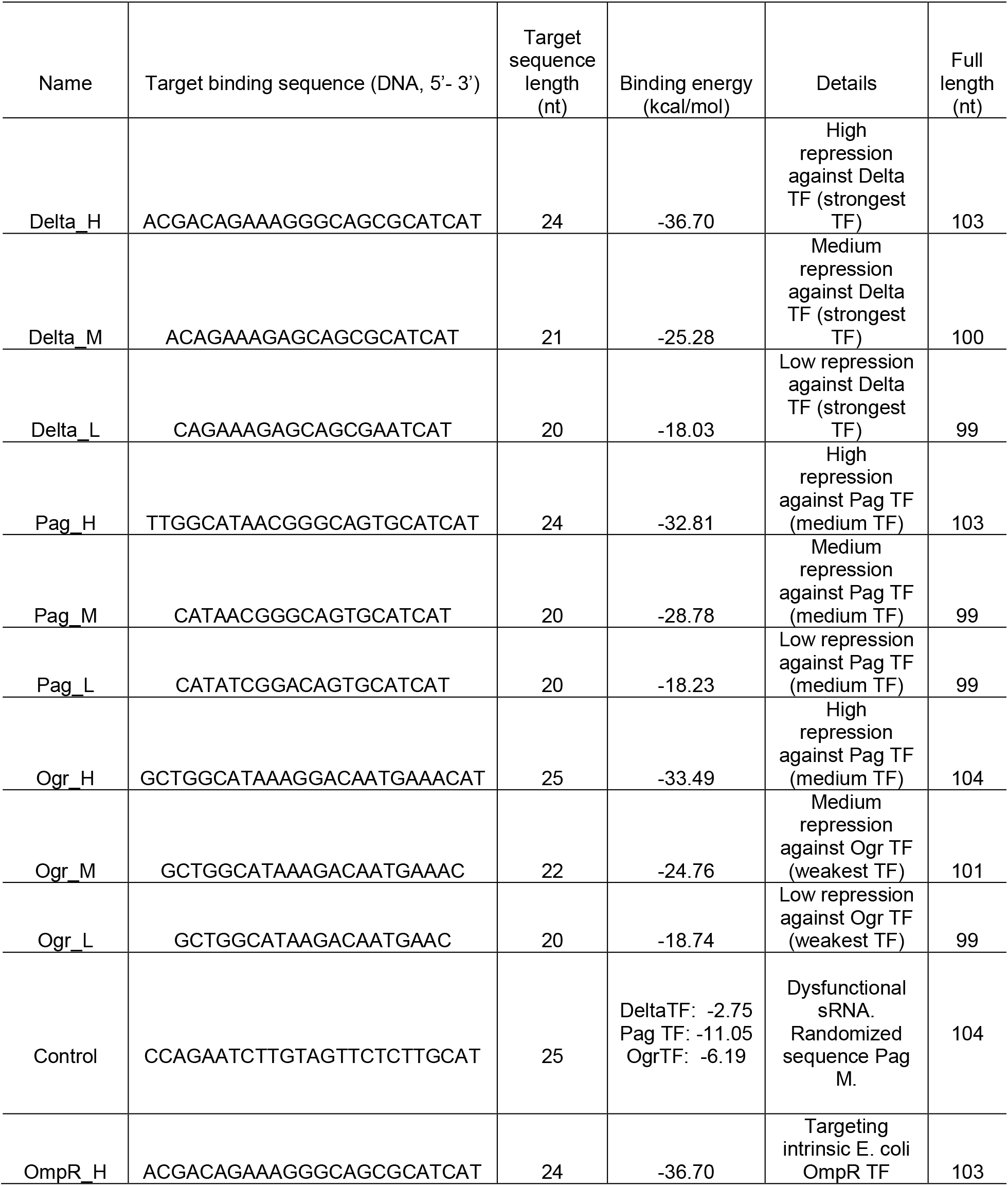
Synthetic sRNAs. Only the target sequence is shown. The 79-nt-long sequence of the MicC scaffold is attached immediately downstream of the target sequence.

**Table S2.** Plasmids used in this study. pGGA is a high copy number plasmid with chloramphenicol resistance; it has a colE1/pMB1/pBR322 pUC-derived *ori*. pSB3C5 is a low-medium copy plasmid with chloramphenicol resistance; it has a p15A *ori*. pSB1A3 is a high copy plasmid with ampicillin resistance; it has a with pUC19-derived pMB1 *ori*.

| Plasmid name | Backbone/Vector | Length (bp) | Construct |
| --- | --- | --- | --- |
| **Genetic circuit plasmids** |  |  |  |
| G1_delta | pGGA | 3641 | pBAD promoter -- (Delta TF) -- *rrnB* T1 terminator -- GFP -- pF promoter |
| G1_pag | pGGA | 3614 | pBAD promoter -- (Pag TF) -- *rrnB* T1 terminator -- GFP -- pF promoter |
| G1_ogr | pGGA | 3614 | pBAD promoter -- (Ogr TF) -- *rrnB* T1 terminator -- GFP -- pF promoter |
| G2_delta | pSB3C5 | 4279 | pBAD promoter -- (Delta TF) -- *rrnB* T1 terminator -- GFP -- pF promoter |
| G2_pag | pSB3C5 | 4252 | pBAD promoter -- (Pag TF) -- *rrnB* T1 terminator -- GFP -- pF promoter |
| G2_ogr | pSB3C5 | 4252 | pBAD promoter -- (Ogr TF) -- *rrnB* T1 terminator -- GFP -- pF promoter |
| G3_delta | pSB3C5 | 5393 | pBAD promoter -- (Delta TF) -- *rrnB* T1 terminator -- GFP -- pF promoter - AraC |
| G3_pag | pSB3C5 | 5503 | pBAD promoter -- (Pag TF) -- *rrnB* T1 terminator -- GFP -- pF promoter - AraC |
| G3_ogr | pSB3C5 | 5477 | pBAD promoter -- (Ogr TF) -- *rrnB* T1 terminator -- GFP -- pF promoter - AraC |
| BBa_K1909013<sup>a</sup> | pSB1C3 | 2070 | pOmpR-GFP-terminator |
| **sRNA-expressing plasmids** |  |  |  |
| sRNA_Delta_H | pSB1A3 | 2275 | T7 promoter - Delta_H sRNA - mic C scaffold |
| sRNA_Delta_M | pSB1A3 | 2272 | T7 promoter - Delta_M sRNA - mic C scaffold |
| sRNA_Delta_L | pSB1A3 | 2271 | T7 promoter - Delta_M sRNA - mic C scaffold |
| sRNA_Pag_H | pSB1A3 | 2275 | T7 promoter - Pag_H sRNA - mic C scaffold |
| sRNA_Pag_M | pSB1A3 | 2271 | T7 promoter - Pag_H sRNA - mic C scaffold |
| sRNA_Pag_L | pSB1A3 | 2271 | T7 promoter - Pag_H sRNA - mic C scaffold |
| sRNA_Ogr_H | pSB1A3 | 2276 | T7 promoter - Ogr_H sRNA - mic C scaffold |
| sRNA_Ogr_M | pSB1A3 | 2273 | T7 promoter - Ogr_M sRNA - mic C scaffold |
| sRNA_Ogr_L | pSB1A3 | 2271 | T7 promoter - Ogr_H sRNA - mic C scaffold |
| sRNA_sRNA_C | pSB1A3 | 2271 | T7 promoter -- randomized sRNA - mic C scaffold |

<sup>a</sup>pSB1C3 BBa\_K1909013 plasmid was a kind gift from the iGEM team Hamburg 2016.

**Table S3.**
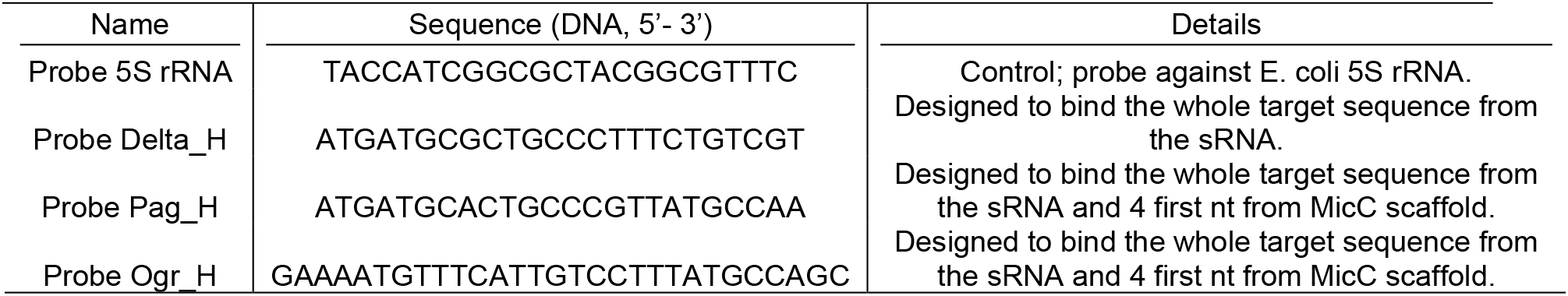
Sequences of the DNA probes used in the northern dot blots to detect high repression sRNAs for each TF. All probes were Cy3 labeled.

**Table S4.**
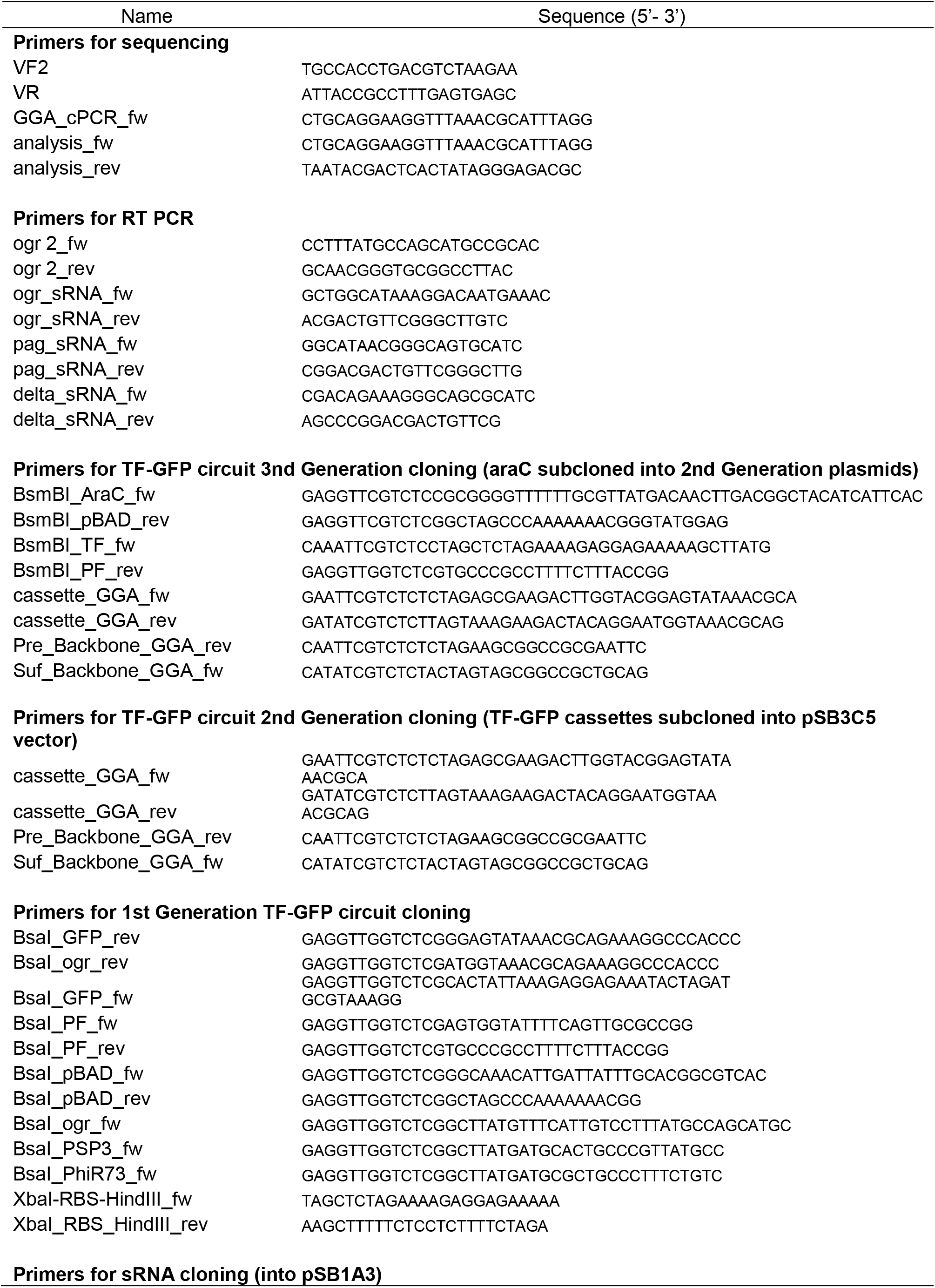
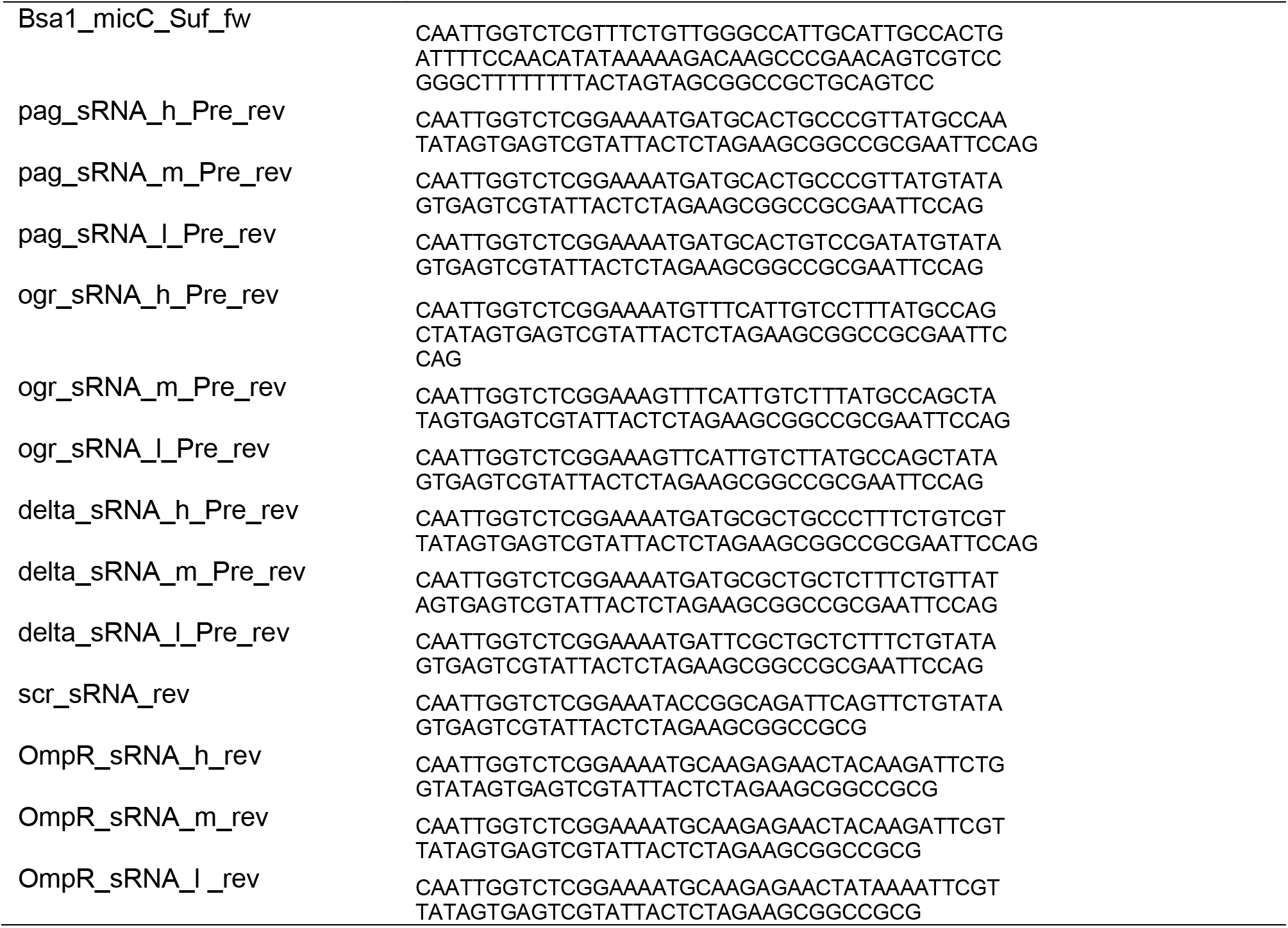
Oligonucleotide primers used in this study. fw, forward primer, rev, reverse primer.

**Table S5.**
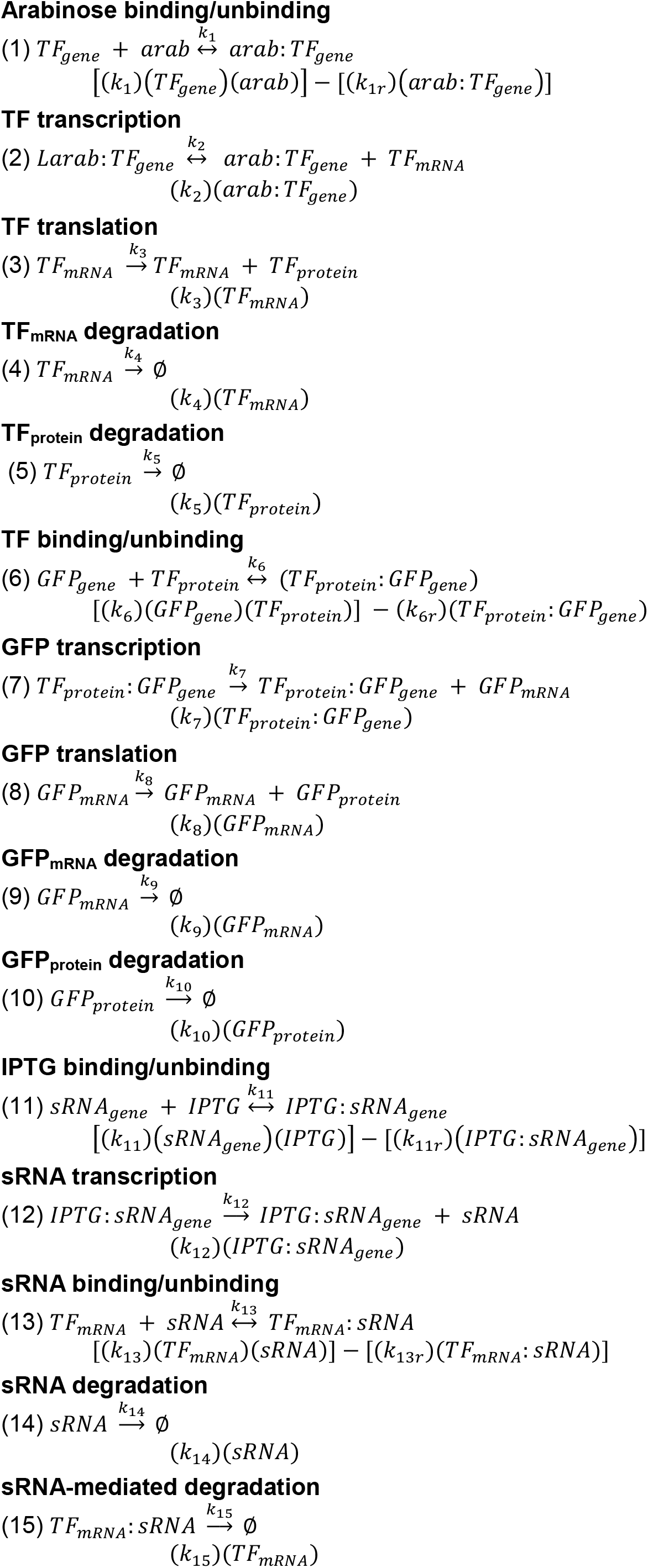
Kinetic model based on the chemical reaction network (CRN). Equation indexes are in accordance to the diagram on Figure S8. Molecular duplexes (complexes) are represented with (:)

**Table S6.** Parameters and simulation details. Deterministic simulations were performed as time-course plots for 10 minutes with following parameters.

| Name | Type | Scope | Initial Value | Units |
| --- | --- | --- | --- | --- |
| cell | compartment | cell | 1 | liter |
| TF<sub>gene</sub> | species | cell | 500 | molecule |
| Larab:TF<sub>gene</sub> | species | cell | 0 | molecule |
| TF<sub>mRNA</sub> | species | cell | 0 | molecule |
| TF<sub>protein</sub> | species | cell | 0 | molecule |
| Larab | species | cell | 500 | molecule |
| GFP<sub>gene</sub> | species | cell | 0 | molecule |
| STF<sub>protein</sub>:GFP<sub>gene</sub> | species | cell | 500 | molecule |
| GFP<sub>mRNA</sub> | species | cell | 0 | molecule |
| GFP<sub>protein</sub> | species | cell | 0 | molecule |
| IPTG | species | cell | 2000 | molecule |
| sRNA<sub>gene</sub> | species | cell | 2000 | molecule |
| IPTG:sRNA<sub>gene</sub> | species | cell | 0 | molecule |
| sRNA | species | cell | 0 | molecule |
| TF<sub>mRNA</sub>:sRNA | species | cell | 0 | molecule |
| k<sub>1</sub> | parameter | arabinose binding | 0.2 | 1/(second*molecule) |
| k<sub>1r</sub> | parameter | arabinose unbinding | 1 | 1/second |
| k<sub>2</sub> | parameter | cell | 0.2 | 1/second |
| k<sub>3</sub> | parameter | TF translation | 20 | 1/second |
| k<sub>4</sub> | parameter | TF<sub>mRNA</sub> degradation | 1.5 | 1/second |
| k<sub>5</sub> | parameter | TF<sub>protein</sub> degradation | 1 | 1/second |
| k<sub>6</sub> | parameter | TF binding | 0.2 | 1/(molecule*second) |
| k<sub>6r</sub> | parameter | TF unbinding | 1 | 1/second |
| k<sub>7</sub> | parameter | cell | 0.3 | 1/second |
| k<sub>8</sub> | parameter | GFP translation | 20 | 1/second |
| k<sub>9</sub> | parameter | GFP<sub>mRNA</sub> degradation | 1.5 | 1/second |
| k<sub>10</sub> | parameter | GFP<sub>protein</sub> degradation | 1 | 1/second |
| k<sub>11</sub> | parameter | IPTG binding | 0.2 | 1/(second*molecule) |
| k<sub>11r</sub> | parameter | IPTG unbinding | 1 | 1/second |
| k<sub>12</sub> | parameter | sRNA transcription | 0.8 | 1/second |
| k<sub>13</sub> | parameter | sRNA:TF<sub>mRNA</sub> binding | 0.5 | 1/(second*molecule) |
| k<sub>13r</sub> | parameter | sRNA:TF<sub>mRNA</sub> unbinding | 1 | 1/second |
| k<sub>14</sub> | parameter | sRNA degradation | 1.5 | 1/second |
| k<sub>15</sub> | parameter | sRNA-mediated degradation | 5 | 1/second |

## Notes

### Competing Interest Statement

The authors have declared no competing interest.

